# LBH is a cancer stem cell- and metastasis-promoting oncogene essential for WNT stem cell function in breast cancer

**DOI:** 10.1101/2021.01.29.428659

**Authors:** Koteswararao Garikapati, Kilan Ashad-Bishop, Sunhwa Hong, Rehana Qureshi, Megan E. Rieger, Linsey E. Lindley, Bin Wang, Diana J. Azzam, Mahsa Khanlari, Mehrad Nadji, Chaitanya Jain, Deukwoo Kwon, Yuguang Ban, Zhen Gao, Steven X. Chen, Andrew H. Sims, Susan B. Kesmodel, Joyce M. Slingerland, Karoline J. Briegel

## Abstract

Cancer stem cells (CSCs) initiate tumors, resist treatment, and seed lethal metastases; yet CSC-specific treatments are lacking. Aggressive, treatment-resistant triple-negative breast cancers (TNBC) exhibit WNT pathway activation and are CSC enriched. Here, we show that Limb-Bud-and-Heart (LBH), a WNT/β-catenin target required for normal mammary stem cell self-renewal, marks poor prognosis, stem-like TNBC, and is a key controller of breast cancer stemness. LBH is specifically expressed in tumor-initiating CD44^+^CD24^-/low^ breast CSCs. LBH overexpression confers stem-like, metastatic traits on both TNBC and luminal origin, non-TNBC breast cancer cells by activating stem cell transcriptional programs. Importantly, silencing LBH potently suppresses tumor initiation and metastasis *in vivo*, and sensitizes TNBC cells to chemotherapy. LBH knockout in the MMTV-Wnt1 breast cancer mouse model, furthermore, revealed LBH is required for WNT-driven breast CSC expansion. Our findings identify LBH as an essential CSC driver downstream of WNT, and a new molecular target for anti-cancer stem cell therapy.

## INTRODUCTION

Triple-negative breast cancers (TNBC) have the worst prognosis of all breast cancer subtypes and contribute disproportionally to cancer death, owing to their innate propensity to metastasize and resist treatment ^1^. Unlike other subtypes of breast cancer (Luminal A/B; HER2+), TNBC lack expression of key targetable markers, Estrogen Receptor (ER), Progesterone Receptor (PR), and/or Human Epidermal Growth Factor Receptor 2 (HER2) ^1^. Consequently, the mainstay of therapy remains chemotherapy, to which >50% of TNBC patients exhibit either *de novo* resistance or rapidly acquire it during an aggressive disease course, often culminating in death within 5 years of diagnosis ^1^. Thus, novel TNBC-specific biomarkers and molecular targets are needed.

TNBC are a heterogeneous group of mostly high-grade (undifferentiated) cancers, of which >80% exhibit immunopositivity for basal cytokeratin 5/6 (K5/6) ^1,2^, a marker usually found in basal/myoepithelial cells of the normal breast. Intriguingly, basal-like TNBC show frequent hyperactivation of stem cell signaling pathways, including the WNT ^3,4^ and TGFβ pathways ^5^, and greater enrichment for malignant cancer stem/progenitor cells than observed in other breast cancer types ^6,7^. Cancer stem cells (CSCs) are a subset of stem-like tumor cells expressing specific markers (CD44^+^CD24^-/low^; ALDH1^+^) and gene signatures of normal mammary stem/progenitor cells ^8,9^. CSCs have increased self-renewal capacity *in vitro* and tumor initiating and metastatic potential *in vivo* ^8,9^, as well as resistance to chemotherapy and radiation ^10,11^. They are, thus, critical mediators not only of tumor initiation, but also of treatment resistance, cancer recurrence, and the main cause of cancer death through metastasis. While it is thought that CSC are responsible for the innate aggressiveness of TNBC ^12^, the mechanisms sustaining high CSC abundance in TNBC remain poorly understood. Furthermore, although the outgrowth of treatment-resistant CSCs represents a serious clinical problem, no CSC-specific therapies have yet been approved in the clinic.

We previously identified Limb-Bud-and-Heart (LBH), a highly conserved, tissue-specific transcription co-factor in vertebrates ^13,14^ that forms its own protein class ^15^. We showed *LBH* is a direct WNT/β-catenin target gene, widely induced by canonical WNT signaling in epithelial tissues ^16^, including in breast epithelial and in TNBC cells ^16,17^. LBH is also a TGFβ/SMAD3 target gene, however, induction of LBH by TGFβ appears context specific ^18^. In normal mammopoiesis, LBH is expressed in multi-potent mammary stem cells (MaSCs) located in the basal epithelium of the postnatal mammary gland, including in WNT-responsive LGR5+ MaSCs ^19^. *Lbh* knockout studies in mice have demonstrated LBH function is required for MaSC self-renewal and differentiation underlying mammary gland outgrowth ^19^. Notably, LBH is upregulated in mammary tumors of MMTV-WNT1^Tg^ mice ^16,20^, and *Lbh* knockout in this WNT-driven breast cancer mouse model attenuated mammary hyperplasia and tumor onset ^20^. Significantly, *in silico* analysis of published cancer gene expression data has suggested that in human breast cancer, *LBH* is most prevalently overexpressed in basal-like TNBC tumors, in association with WNT pathway activation ^16^.

Overexpression of LBH has also been described in gastric cancers and glioblastoma, where it promotes cell proliferation, invasion, angiogenesis, and tumor growth ^21,22^. In contrast, LBH is downregulated in nasopharyngeal and lung cancer, where it has non-invasive, tumor suppressive effects by inducing G1/S cell cycle arrest ^23,24^. However, the clinical and functional relevance of aberrant LBH overexpression in breast cancer is still poorly understood. Moreover, while LBH appears critical to normal MaSC regulation ^19^, and for WNT-induced mammary tumorigenesis ^20^, it’s potential role in WNT-mediated stem cell control has not been characterized. Through comprehensive analysis of LBH protein expression and function in multiple, tumor-subtype specific primary and established breast cancer cell models, clinical breast cancer specimen, genetic mouse models expressing stem cell reporters, and RNA-Seq, we here uncovered a novel role of LBH as a CSC-driving oncogene that is crucial for TNBC metastasis and chemoresistance, as well as WNT-driven breast CSC amplification *in vivo.* Importantly, silencing LBH attenuates tumor initiation and metastasis, while sensitizing TNBC cells to chemotherapy. Mechanistically, LBH activates mammary stem cell transcriptional programs and represses epithelial adherence junction genes. Unlike other CSC-promoting factors, however, LBH does not induce mesenchymal traits. Thus, LBH is a unique CSC-driver downstream of WNT, and a new molecular target for CSC-specific cancer therapy.

## RESULTS

### LBH expression in human breast cancer strongly associates with TNBC, and expression of stem cell and metastatic gene profiles

To date there has been no analysis of LBH protein expression in clinical breast cancer samples; thus, we first performed immunohistochemical (IHC) analysis of 250 newly diagnosed invasive primary breast cancers (Table 1; and representative IHC images shown in Fig. 1a). High LBH protein expression (defined as immunostaining intensity of >+2 and >50% nuclei positive) was detected in 16% of all breast tumors (40/250; p<0.001; Table 1). High LBH most strongly correlated with triple-negative (ER-/PR-/HER2-) status, as 55% of LBH-positive tumors were TNBC (p=0.0001; Table 1). In TNBC tumors, LBH was overexpressed in both tumor nuclei and cytoplasm compared to its nuclear, basal cell-restricted expression in normal breast tissues (Fig. 1a) ^19^. In contrast, LBH was negatively correlated with hormone receptor (ER and PR) expression (p=0.0002 for ER-, and p=0.0030 for PR-), a feature of luminal breast cancers, and HER2+ status (p=0.0174; Table 1) (Fig. 1a). Notably, LBH strongly associated with high tumor grade (III) (p=0.0092), but not with TNM stage (p=0.3128) or lymph node status (p=0.4978) in this breast cancer cohort (Table 1).

**Figure 1:**
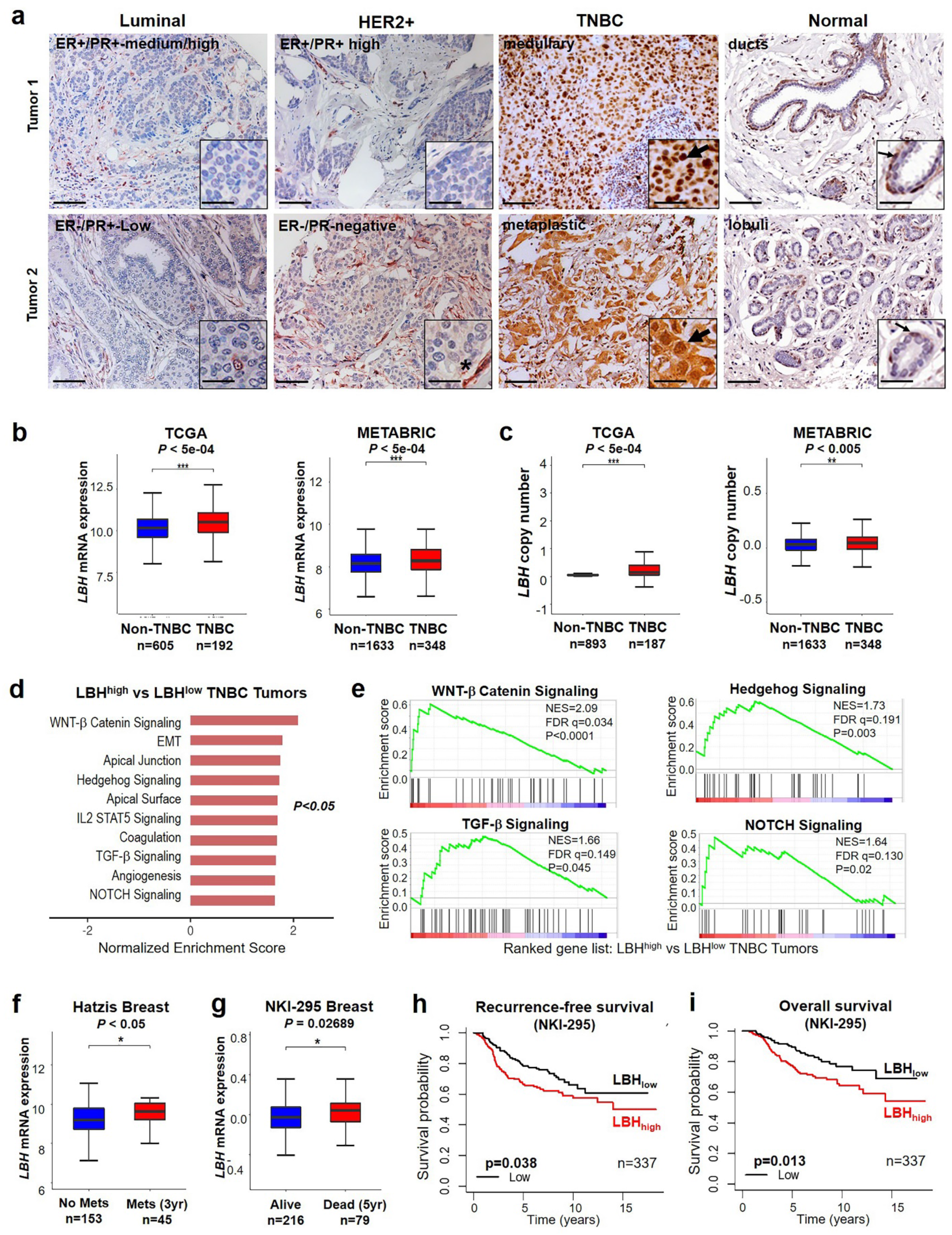
LBH expression in human breast cancer associates with TNBC, stemness, metastasis, and poor outcome. (**a**) Representative IHC images of LBH protein expression (brown) in primary, high-grade (II-III) human breast cancers compared to normal breast tissue. Tumor subtypes and hormone receptor (ER, PR) status, as indicated; Scale bars, 100 μm; 50 μm (close-ups). Note, the LBH overexpression (nuclear + cytoplasmic) in TNBC tumor cells (arrow heads) compared to its nuclear, basal cell-specific expression in normal breast epithelium (thin arrows). LBH was also abundant in tumor stroma (asterix) of non-TNBC tumors. (**b,c**) Box plots showing *LBH* mRNA expression (**b**) and copy numbers (**c**) in TNBC (red) versus all other subtypes (non-TNBC; blue) in the TCGA ^25^ (left) and METABRIC ^26^ (right) breast cancer datasets according to their IHC-defined biomarker status. Number (n) of patients per group as indicated. (**d,e**) Gene set enrichment analysis (GSEA): (**d**) Top 10 gene signatures differentially expressed between LBH^high^ (expressing LBH above the median) and LBH^low^ TNBC in the TCGA data set (p<0.05 and FDR<0.25); (**e**) Enrichment plots of stem cell signaling pathways in LBH^high^ vs. LBH^low^ TNBC. NES, Normalized Enrichment Score; FDR, False Discovery Rate. (**f,g**) Correlation of high *LBH* expression (top quartile; red) with (**f**) metastatic event at 3 years (yr) and (**g**) patient death at 5 years after diagnosis. Gene expression data were obtained from the Hatzis ^62^ and NKI-295 ^61^ BrCa cohorts. (**h,i**) Kaplan-Meier plots showing *greater-than-median* intra-tumor levels of *LBH* (LBHhigh, red) associate with: (**h**) reduced recurrence-free survival and (**i**) overall survival in the NKI-295 cohort. Data represent means ± SEM. P-values, as indicated; *P<0.05; **P<0.01; ***P<5e-04. Mann-Whitney U test (**b-c, f-g**), and log rank test (**h-j**) was used.

**Table 1.** Association of LBH expression with clinical parameters in breast cancer patients

| Variable | All |  | LBH<br>Low |  | LBH<br>High |  | *p-value*<sup>#</sup> |
| --- | --- | --- | --- | --- | --- | --- | --- |
|  | *n* | % | *n* | % | *n* | % |  |
| **Total number of IDCs** | **250** | **100** | **210** | **84.0** | **40** | **16.0** | **<.0001<sup>\$</sup>** |
| Estrogen Receptor |  |  |  |  |  |  | **0.0002** |
| *Negative* | 145 | 58.0 | 111 | 52.9 | 34 | 85.0 |  |
| *Positive* | 105 | 42.0 | 99 | 47.1 | 6 | 15.0 |  |
| Progesterone Receptor |  |  |  |  |  |  | **0.0030** |
| *Negative* | 154 | 61.6 | 121 | 57.6 | 33 | 82.5 |  |
| *Positive* | 96 | 38.4 | 89 | 42.4 | 7 | 17.5 |  |
| HER2 |  |  |  |  |  |  | **0.0174** |
| *Negative* | 132 | 52.8 | 104 | 49.5 | 28 | 70.0 |  |
| *Positive* | 118 | 47.2 | 106 | 50.5 | 12 | 30.0 |  |
| Triple-negative |  |  |  |  |  |  | **0.0001** |
| *Yes* | 74 | 29.6 | 52 | 24.8 | 22 | 55.0 |  |
| *No* | 176 | 70.4 | 158 | 75.2 | 18 | 45.0 |  |
| TNM stage |  |  |  |  |  |  | 0.3128* |
| *Tis or T1 or T2* | 152 | 60.8 | 128 | 61.0 | 24 | 60.0 |  |
| *T3 or T4* | 82 | 32.8 | 73 | 34.8 | 9 | 22.5 |  |
| *Unknown* | 16 | 6.4 | 9 | 4.3 | 7 | 17.5 |  |
| Lymph Node status |  |  |  |  |  |  | 0.4978* |
| *N0 or N1* | 194 | 77.6 | 168 | 80.0 | 26 | 65.0 |  |
| *N2 or N3* | 40 | 16.0 | 33 | 15.7 | 7 | 17.5 |  |
| *Unknown* | 16 | 6.4 | 9 | 4.3 | 7 | 17.5 |  |
| Tumor grade |  |  |  |  |  |  | **0.0092\*** |
| *Low grade* | 31 | 12.4 | 31 | 14.8 | . | . |  |
| *High grade* | 218 | 87.2 | 178 | 84.8 | 40 | 100.0 |  |
| *Unknown* | 1 | 0.4 | 1 | 0.5 | . | . |  |

IDC = Invasive ductal carcinoma. Bold data indicate significant correlation.
<sup>#</sup> Chi-square test was used. <sup>\$</sup> One proportion test was used.
\* Unknown values were excluded.

Bioinformatic interrogation of two independent breast cancer data sets, the TCGA ^25^ and the METABRIC ^26^, that together include over 2,778 primary cancers, further showed *LBH* mRNA (Fig. 1b), and *LBH* gene copy numbers (Fig. 1c), were modestly, but significantly increased in TNBC compared to non-TNBC cancers. PAM50 classification of the METABRIC data into intrinsic molecular breast cancer subtypes ^26^ demonstrated *LBH* mRNA levels were highest in basal, normal-like, and rare claudin-low (Supplementary Fig. 1a), which are enriched in TNBC ^27,28^, and lowest in luminal-type breast cancers, consistent with our IHC data and previous gene profiling data ^16,18^. Notably, when grouped into ten prognostic Integrative Clusters (IntClust = IC), *LBH* expression was greater in TNBC clusters (IC4ER-; IC10, basal; Supplementary Fig. 1b), with the highest risk of disease relapse and death ^29^. Finally, subclassification of primary TNBC into four molecular subgroups, basal-like (BL1, BL2), mesenchymal (M), and luminal androgen receptor (AR) positive (LAR), according to Lehman et al. ^2,30^ revealed higher *LBH* mRNA levels in aggressive basal-like and mesenchymal TNBC tumors compared to the more indolent LAR subgroup (Supplementary Fig. 1c). Thus, LBH is upregulated in TNBC at the protein, RNA, and gene expression levels.

To identify pathways associated with LBH in clinical TNBC, we performed Gene Set Enrichment Analysis (GSEA) of primary TNBC tumors from the TCGA dataset (Fig. 1b). Remarkably, 4 of the top 10 significantly enriched signatures in LBH^high^ TNBC were stem cell signaling pathways (WNT, Hedgehog, TGFβ, Notch). WNT/β-catenin signaling genes were most significantly enriched (NES = 2.09, p<0.0001, q=0.034; Fig. 1d,e), consistent with our previous finding that *LBH* is a WNT target gene ^16^. LBH^high^ TNBC tumors also expressed an EMT gene signature (Fig. 1d), indicating *LBH* expression in TNBC associates with both stem cell and metastatic gene profiles.

To further investigate the clinical relevance of LBH overexpression in breast cancer, we analyzed *LBH* gene expression in cohorts, representative of the patient population as a whole. Importantly, high intra-tumoral *LBH* expression (top quartile) in primary human breast cancers significantly associated with advanced disease stage (p=0.001; Supplementary Fig. 1d), as well as metastatic event (p<0.05; Fig. 1f) and early patient death 3-5 years after diagnosis (p=0.027; Fig. 1g). Moreover, patients with LBH^high^ cancers (expressing *LBH* above the median) had reduced relapse-free [RFS] (p=0.038) and overall survival [OS] (p=0.013) compared to those with LBH^low^ tumors (Fig. 1h,i). Collectively, these data demonstrate that LBH is a putative biomarker for TNBC, and its expression might herald the presence of stem-like, metastatic cancer cells.

### LBH is required for clonogenicity and *in vivo* tumor growth of TNBC cells

LBH protein expression in human breast cancer cell lines showed a similar pattern. It was observed exclusively in TNBC, and not detected in 7 different non-TNBC-derived breast cancer lines (Fig. 2a,b). Moreover, nearly all LBH-positive TNBC lines (6/7) expressed high levels of CD44, a marker of breast CSCs (Fig. 2a). To investigate the functional relevance of LBH overexpression in TNBC, we depleted LBH in both, basal-like and mesenchymal TNBC cell line models with high endogenous LBH expression, HCC1395 and MDA-MB-231, respectively (Fig. 2a,c) ^16^. Immediate effects of LBH on tumorigenicity *in vitro* were assayed by comparing transient cell transfection of small inhibitory RNAs (siRNA) targeting LBH (siLBH) with non-targeted siRNA (siCtrl) controls. LBH knockdown in HCC1395 and MDA-MB-231, as confirmed by qPCR and Western blot analyses (Fig. 2c), significantly attenuated the increase in cell numbers over time (Fig. 2d) and diminished clonogenicity in 2D culture (Fig. 2e), as well as anchorage-independent growth in soft agar (Fig. 2f), a key feature of malignant transformation.

**Figure 2.**
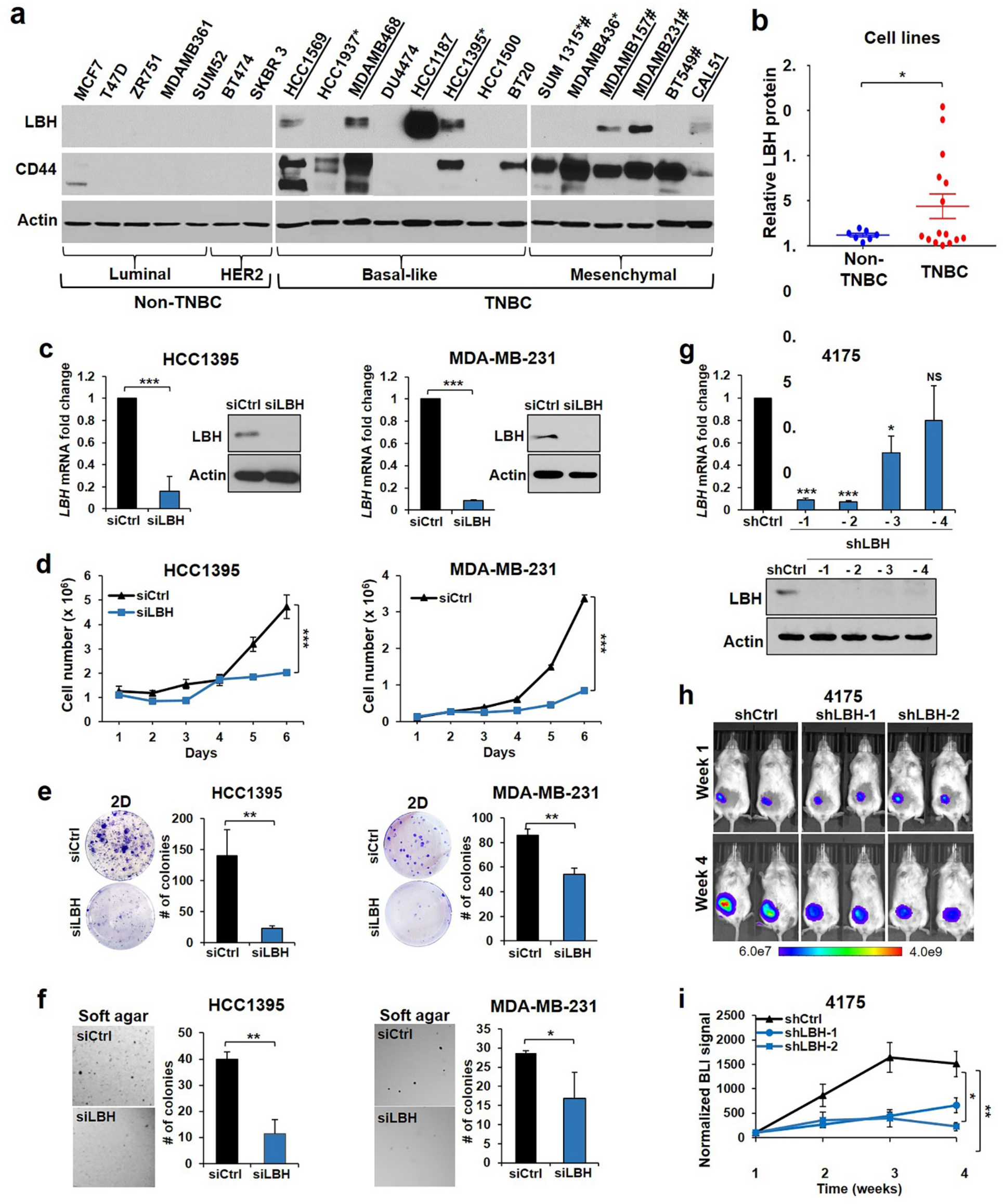
LBH is required for clonogenicity and *in vivo* tumor growth of TNBC cells. (**a**) Western blot analysis of LBH and stem cell marker, CD44, in human breast cancer cell lines. Subtypes, LBH expression (underlined), BRCA1 mutant (asterisk) and claudin-low (number sign) status, as indicated. See also Supplementary Fig. 1. (**b**) Relative LBH protein in TNBC (n=14) *vs.* non-TNBC lines (n=7) quantified by densitometry relative to β-actin loading control. \**P*<0.05, Twotailed Student’s *t*-test. (**c-f**) TNBC cell lines, HCC1395, MDA-MB-231, were transiently transfected with a pool of four independent LBH-targeted siRNAs (siLBH), or scrambled siRNA control (siCtrl). (**c**) qPCR (left) and Western blot (right) showing LBH expression 3 or 4 days after siRNA transfection, respectively. (**d**) Proliferation (cell number over time) in adherent cell cultures normalized to values at day 0. (**e**) Two-dimensional (2D) colony formation, and (**f**) anchorage-independent growth in soft agar. Representative images (left) and quantification of colony numbers (#) (right). Experiments were repeated > three times with n = 3 replicates per group. (**g, h**) Luciferase-tagged MDA-MBA-231 variant, 4175, was stably transduced with non-target control shRNA (shCtrl) or four different *LBH*-targeted shRNAs (shLBH1-4). (**g**) qPCR (left) and Western blot (right) show efficient LBH knockdown (>85%) in 4175 TNBC cells transduced with shLBH-1 and shLBH-2. (**h**) Representative bioluminescence (BLI) images after orthotopic mammary fat pad injection of 4175 cells (10^5^) expressing shCtrl, shLBH-1, or shLBH-2 into female NSG mice. (**i**) Quantification of tumor growth by weekly BLI measurements, normalized to BLI signals (photons per second) at day 1. This experiment was repeated twice with n = 6 mice per group each. All data represent means ± SEM. \**P* < 0.05, \*\**P* < 0.01, \*\*\**P* < 0.001. Two-tailed Student’s *t*-test (**b-c, e-f**), or non-linear regression models (**d,i**) were used.

To test the effects of LBH on tumorigenicity of TNBC cells *in vivo*, a luciferase-positive subclone of MDA-MB-231, 4175 ^31^, was stably transduced with either non-target shRNA control (shCtrl), or with one of four independent LBH-targeted shRNAs (Fig. 2g). Two different LBH shRNA transduced 4175 populations (shLBH-1, shLBH-2) with the greatest LBH knockdown efficiencies (85-90%; Fig. 2g), or shCtrl transduced cells were implanted into the mammary fat pads of immunocompromised female *NOD/SCID-Il2Rgamma-/-* (NSG) mice (10^5^ cells/mouse; 12 mice/group). Whole animal *in vivo* Bioluminescence Imaging (BLI) analysis revealed that LBH KD significantly attenuated *in vivo* tumor growth (Fig. 2h,i).

### High LBH levels promote breast cancer growth *in vivo*

We next examined whether LBH overexpression would increase breast tumor development. *LBH* was transfected stably into BT549, a mesenchymal TNBC line with low to undetectable endogenous LBH (Fig. 2a and Fig. 3a) ^16^. To test if oncogenic effects of LBH are TNBC-specific, *LBH* was also introduced into MCF7, an ER+ luminal (non-TNBC) breast cancer cell model, lacking LBH expression (Fig. 2a and Fig. 3a) ^16^. Ectopic LBH expression in both BT549 and MCF7 yielded LBH expression levels comparable to those in LBH-positive TNBC lines (Fig. 2a and Fig. 3a). While LBH overexpression had little or no effect on cell numbers over 6 days in 2D culture (Fig. 3b), it significantly and consistently increased anchorage-independent growth of both BT549 and MCF7 (Fig. 3c). Importantly, orthotopic injection of luciferase tagged MCF7+LBH into NSG mice (2×10^6^, n=9/group) showed ectopic LBH increased *in vivo* tumor formation (Fig. 3d), tumor volumes (Fig. 3e,f), weight and size (Fig. 3g) compared to MCF7+vector controls, demonstrating LBH is sufficient to increase the breast cancer tumorigenicity. When near-to-limiting dilutions of cells were injected orthotopically, the differences in volumes between MCF7+LBH and MCF7+vector derived tumors became even more notable (p<0.05; Fig. 3h). Thus, LBH is not only required for TNBC tumorigenesis, but also promotes *in vivo* growth in this non-TNBC model, supporting the notion that LBH is a potent breast cancer promoter.

**Figure 3.**
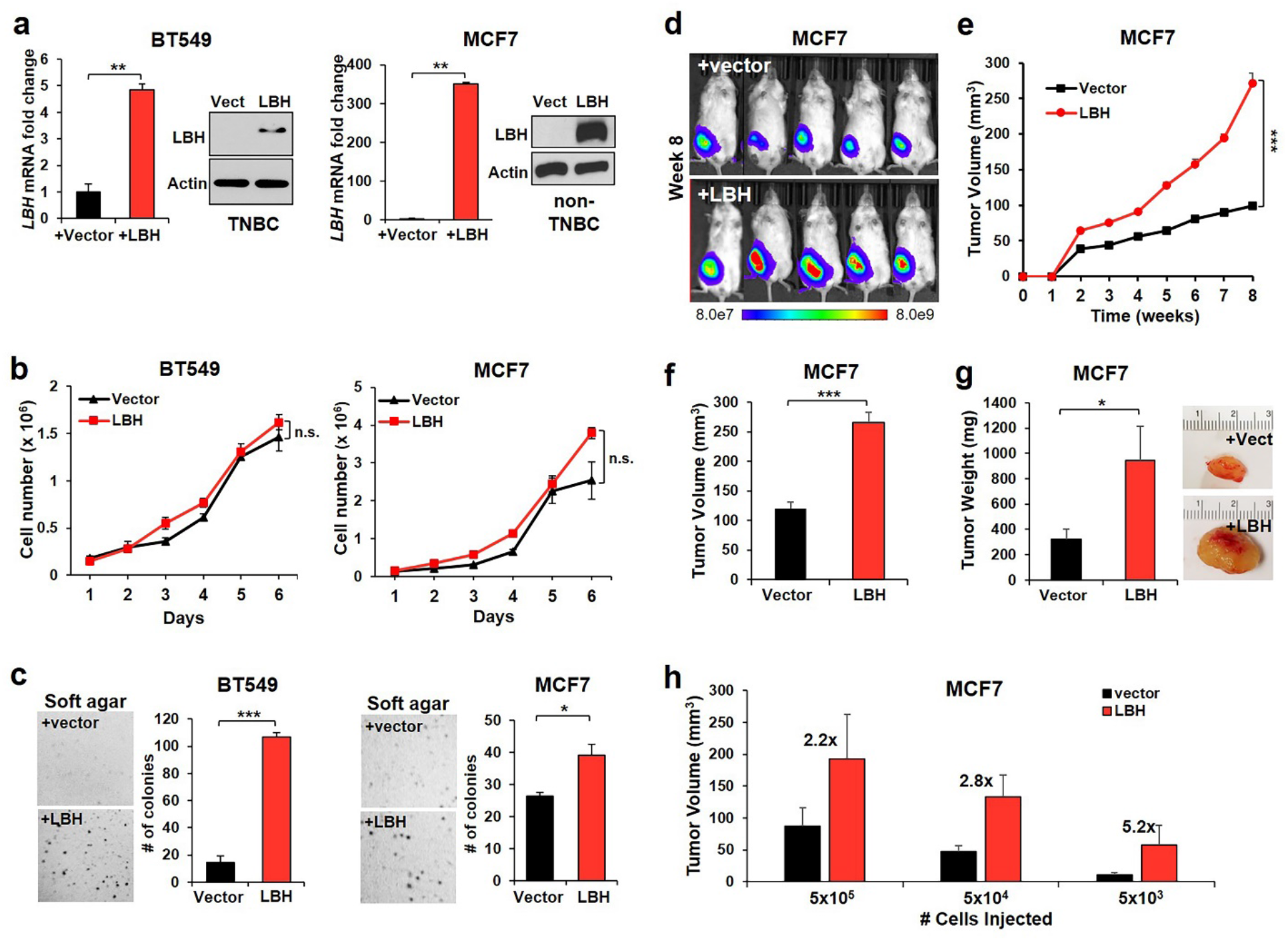
LBH promotes tumorigenicity of TNBC and non-TNBC cells. (**a-c**) BT549 (TNBC) and MCF7 (non-TNBC) BrCa cell lines stably transfected with pCDNA3-LBH expression plasmid (+LBH) or empty vector control (+vector). (**a**) qPCR (left) and Western blot (right) show ectopic LBH expression in BT549+LBH and MCF7+LBH cells. (**b**) Proliferation (cell number over time) in adherent cell cultures (normalized to day 0). (**c**) Soft agar colony formation assay. Representative images (right) and quantification of colony numbers (#) (left). Experiments were repeated > three times with n = 3 replicates per group. (**d-g**) Luciferase-tagged MCF+LBH and MCF7+vector control cells (2 x 10^6^) were injected into the fourth mammary fat pad of NSG hosts. (**d**) Representative BLI images of *in vivo* tumor growth. (**e**) Tumor volumes over time. Vector group (n=9) and LBH group (n=8). (**f**) Tumor volumes and (**g**) Tumor weights at endpoint. Vector group (n=9) and LBH group (n=3). Representative tumor images (right). This experiment was repeated twice with n=6-9 mice per group each. (**h**) Tumor volumes at 12 weeks post orthotopic injection of reducing concentrations of MCF+LBH or MCF7+vector cells into NSG hosts. n=6-8 mice per group (*P* < 0.05). Data represent means ± SEM. \**P* < 0.05; \*\**P* < 0.01, \*\*\**P* < 0.001. n.s., not significant. Two-tailed Student’s *t*-test (**a,c,f,g**), non-linear regression models (**b,e**), or Wilcoxon signed rank test (**h**) were used.

### LBH is enriched in breast CSCs and sufficient to increase CSC abundance

Since LBH controls stem cells in the normal breast ^19^, we next considered the possibility that LBH might promote breast tumorigenesis by modulating CSCs, or tumor-initiating stem cells (TIC). To this end, we first performed serial mammosphere assays, a quantitative *in vitro* measurement of stem cell self-renewal ^32^. Depletion of LBH in four independent TNBC lines HCC1395, MDA-MB-231, as well as HCC1187, a basal *BRCA1^WT^*, and MDA-MB-157, a metaplastic TNBC model, significantly reduced primary (Fig. 4a; and Supplementary Fig. 2a,b), and to a greater extent, secondary sphere formation (2.7 fold in HCC1395/p=0.00176; 2.9 fold in MDA-MB-231/p=0.00154; Fig. 4a). Importantly, LBH overexpression in both BT549, and MCF7 increased primary, and more notably secondary tumor sphere formation (~3.5 fold for BT549+LBH/p<0.001; ~2 fold for MCF7+LBH/p=0.00024) *vs.* the respective controls (Fig. 4b). These results suggest, LBH may augment TI-SC abundance by promoting self-renewal.

**Figure 4:**
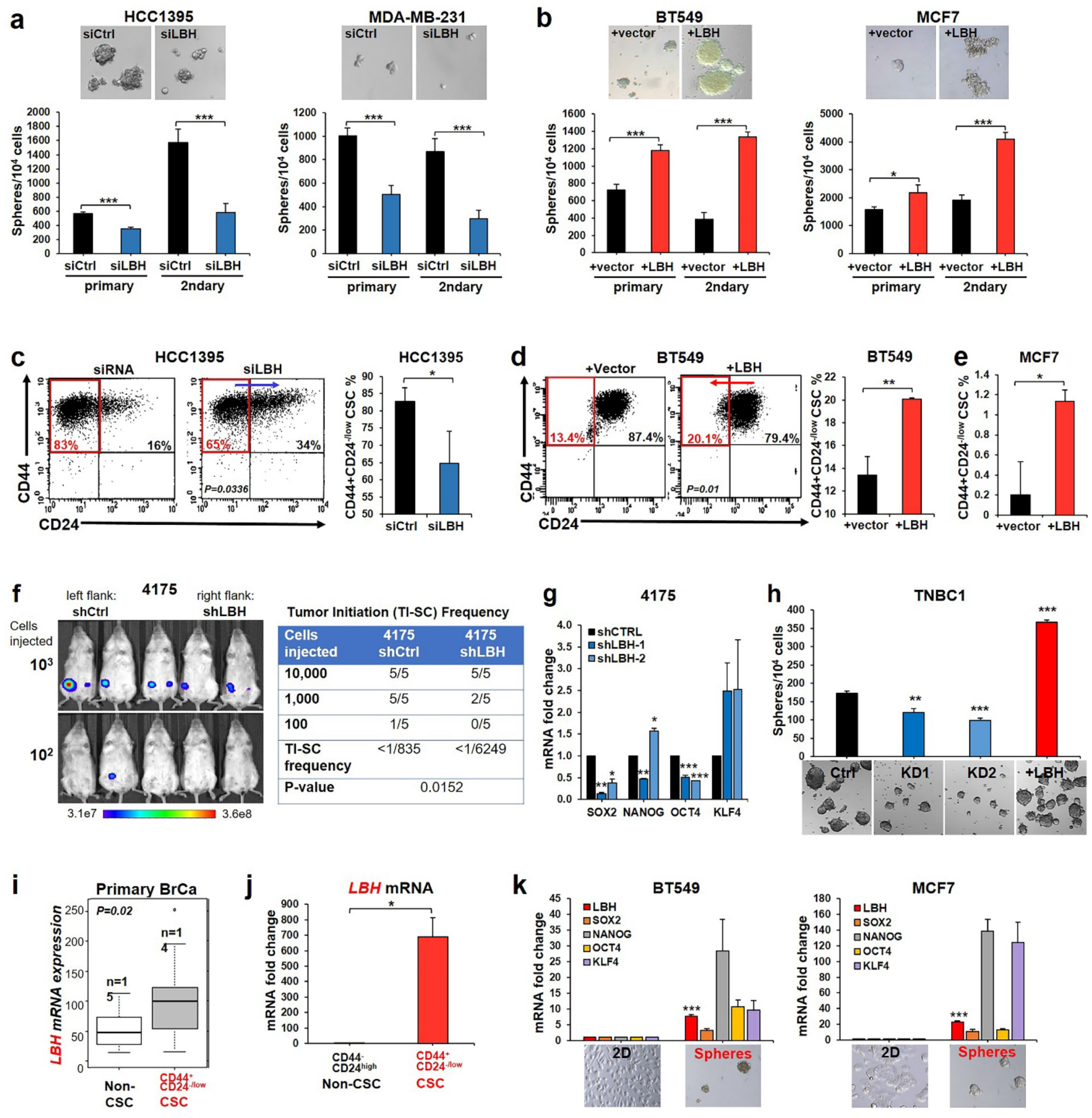
LBH is expressed in breast CSCs and promotes a self-renewing CSC phenotype. (**a,b**) Quantitative serial sphere formation assays with representative tumor sphere images in: (**a**) LBH-depleted HCC1395, MDA-MB-231 TNBC; and (**b**) LBH-overexpressing BT549, MCF7 BrCa cells. (**c-e**) FACS quantification of CD44^+^CD24^-/low^ CSC populations in: (**c**) HCC1395 cells 9 days after transient transfection with siLBH or scrambled siRNA (siCtrl); (**d**) BT549, and (**e**) MCF7 stably expressing LBH (+LBH) or vector control (+vector). Representative FACS plots with gates for CD44^+^CD24^-/low^ CSC (red box) are shown on the right. Blue arrow (in c) indicates increased, red arrow (in d) reduced tumor cell differentiation. (**f**) Limiting cell dilution (10^5^, 10^4^, 10^3^) orthotopic transplant assay of 4175 shCtrl (left flank) and 4175 shLBH-2 (right flank) cells. Representative BLI images at 5 weeks (right) and quantification (Table; left) of tumor-initiating stem cell (TI-SC) frequencies (CI, 95% Confidence Interval; Chi Square Analysis; P=0.0152) using http://bioinf.wehi.edu.au/software/elda/software; n=5 tumors per group. (**g**) qPCR of SC-TFs expression in 4175 shCtrl and LBH shRNA-transduced lines. (**h**) Tumor sphere formation of primary, patient derived TNBC cells (TNBC1) transduced with two independent TET-inducible LBH shRNAs (KD1, KD2), non-target shRNAs (shCtrl), or LBH cDNA (+LBH) expressing lentiviruses in the presence of 1 μg/ml Doxycycline. (**i**) Gene profiling data for *LBH* in a TCGA data set ^36^ comparing gene signatures of CD44^+^CD24^-/low^ CSC (grey bar) and non-CSC (white bar) populations from n>14 primary human breast cancers. *P* value by Mann-Whitney U test. (**j**) qPCR of *LBH* in FACS-sorted CD44^+^CD24^-/low^ CSC (red bar) and CD44^-^CD24^high^ non-CSC (black bar) populations from BrCa lines (see Supplementary Fig. 3). (**c**) qPCR of *LBH* and pluripotency TFs *(SOX2, NANOG, OCT4, KLF4*) expression in CSC-enriched tumor spheres formed by parental BT549 TNBC and MCF7 non-TNBC cells in low-attachment suspension cultures *vs.* cells grown in 2D on adhesive plates. Data (**a-e, g,h,j,k**) represent means ± SEM of three independent experiments. \**P* < 0.05; \*\**P* < 0.01. \*\*\**P* < 0.001 (Student’s t-test; n >3 per group).

To determine whether the increased stem cell activity in the presence of LBH was due to increased CSC abundance, we performed Flow Cytometry Cell Sorting (FACS) analysis, using CD44-CD24 surface markers. We found TNBC lines with endogenous LBH overexpression, HCC1395 and MDA-MB-231, had very high % CD44^+^CD24^-/low^ CSCs (86.3±4.42 and 96.9±0.24 respectively; Supplementary Table 1) compared to the non-LBH expressing BT549 TNBC and MCF7 lines that both had low CD44^+^CD24^-/low^ populations (11.8±1.31 and 0.47±0.31, respectively; Supplementary Table1). Remarkably, LBH depletion in HCC1395, decreased the CD44^+^CD24^-/low^ CSC population by nearly 20% (from 83% to 65%, p=0.03; Fig. 4c), suggesting LBH is required to maintain TI-SCs in TNBC. Conversely, LBH overexpression in BT549 significantly increased the % CD44^+^CD24^-/low^ TI-SC population (from 13.4% to 20.1%; p=0.01, Fig. 4d). Notably, ectopic LBH in MCF7 non-TNBC cells also increased this TI-SC population significantly (p=0.019; Fig. 4d), consistent with the increased tumorigenicity of these cells *in vivo* (Fig. 3d-h).

Interestingly, LBH depletion in MDA-MB-231 and its sister cell line, 4175, did not shift the CD44^+^CD24^-/low^ CSC to the CD44^+^CD24^high^ non-CSC population (Supplementary Fig. 2c), as observed for HCC1395 (Fig. 4c). Rather, in these claudin-low TNBC lines, stable LBH knockdown with two independent shRNAs changed the distribution of CSCs within the CD44^+^CD24^-/low^ subpopulation. LBH depletion caused a five-fold decrease in CD44^+^CD24^low^ CSCs, which are on top of the CSC hierarchy and have the highest self-renewal and metastasis-initiating potential ^33,34^ (from 10.0 to 2.3%; p<0.01; Supplementary Fig. 2c), whereas CD44^+^CD24^-^ cells, with lower TI-SC activity and lacking metastasis-initiating potential ^33,34^, were increased. In concordance with these surface marker changes tumor sphere formation was markedly reduced in MDA-MB-231-shLBH-1/-2 compared to MDA-MB-231-shCtrl control cells (Supplementary Fig. 2a). Thus, LBH shifts surface CD44/CD24 markers towards a more stem-like phenotype and increases the abundance of TI-SCs.^35^

To definitively test the requirement of LBH for self-renewal and TI-SC abundance, we implanted 4175-shCtrl and 4175-shLBH cells at limiting dilutions (10^4^, 10^3^, 10^2^) in either left or right inguinal mammary fat pads of NSG mice. LBH depletion profoundly reduced the TI-SC frequency in 4175 cells (p = 0.0152; Fig. 4f), demonstrating LBH is essential for tumor initiation. Moreover, stem cell transcription factors (SC-TFs), SOX2, OCT4, and NANOG, were downregulated in LBH depleted 4175 lines at both the RNA and protein levels (Fig. 4g; and Supplementary Fig. 2d).

To test if primary breast CSCs also required LBH, we performed LBH knockdown and overexpression studies in a patient derived TNBC cell model (TNBC1) (see Methods). As observed in established cancer lines, in these primary TNBC cells LBH downregulation with two TET-inducible *LBH* shRNAs (KD1, KD2) significantly reduced, whereas transduction with a TET-inducible *LBH* transgene (+LBH) increased tumor sphere formation by > 2-fold (p<0.001; Fig. 4h).

Finally, to test if LBH is expressed in breast CSCs, LBH was evaluated in stem cell enriched and non-stem cell populations of primary human breast cancers. Analysis of published gene signatures of CD44^+^CD24^-/low^ TI-SCs from >14 primary breast cancers ^36^ showed that *LBH* is elevated in this CSC-enriched population compared to non-CSC populations from the same tumors (Fig. 4i). qPCR analysis of FACS-purified stem cell-enriched populations from breast cancer cell lines (Supplementary Fig. 3), furthermore, revealed much higher *LBH* levels in the CD44^+^CD24^-/low^ TI-SC than in the non-TI-SC CD44^-^CD24^high^ population (Fig. 4j).

To further validate the relationship between LBH and cancer stemness, we evaluated the expression of *LBH* and SC-TFs in LBH-low/negative breast cancer cell lines (BT549, MCF7) grown under stem cell enriching conditions in serum-free, low-attachment mammosphere cultures^32^. Notably, *LBH* was co-induced along with key SC-TFs, *SOX2, NANOG, OCT4, KLF4*, in CSC-enriched tumor spheres compared to 2D cultures from both BT549, and MCF7 lines (Fig. 4k). Thus, as for known stem cell drivers, LBH expression increases with stem cell enrichment. Collectively, these data demonstrate that LBH is a novel breast CSC-specific factor necessary and sufficient to promote a malignant CSC phenotype.

### LBH promotes tumor invasion, metastasis, and chemoresistance

CSCs have increased invasive, metastatic properties ^34,37^. Hence, we next explored how LBH affects cell motility and invasion. LBH depletion in TNBC lines (HCC1395, MDA-MB-231, HCC1187, MDA-MB-157) markedly reduced tumor cell migration (Fig. 5a; Supplementary Fig. 4a,b), and invasion (Fig. 5b; Supplementary Fig. 4c). In contrast, LBH overexpression in TNBC and non-TNBC models, BT549 and MCF7 respectively, increased cell motility by >2-4 fold (Fig. 5c), and invasion by BT549 over 5-fold (Fig. 5d). Additionally, whereas control BT549 cells formed small compact spheres in three-dimensional (3D) Matrigel cultures, LBH expressing BT549+LBH formed spheroids with extensive, highly invasive protrusions (Fig. 5e).

**Figure 5.**
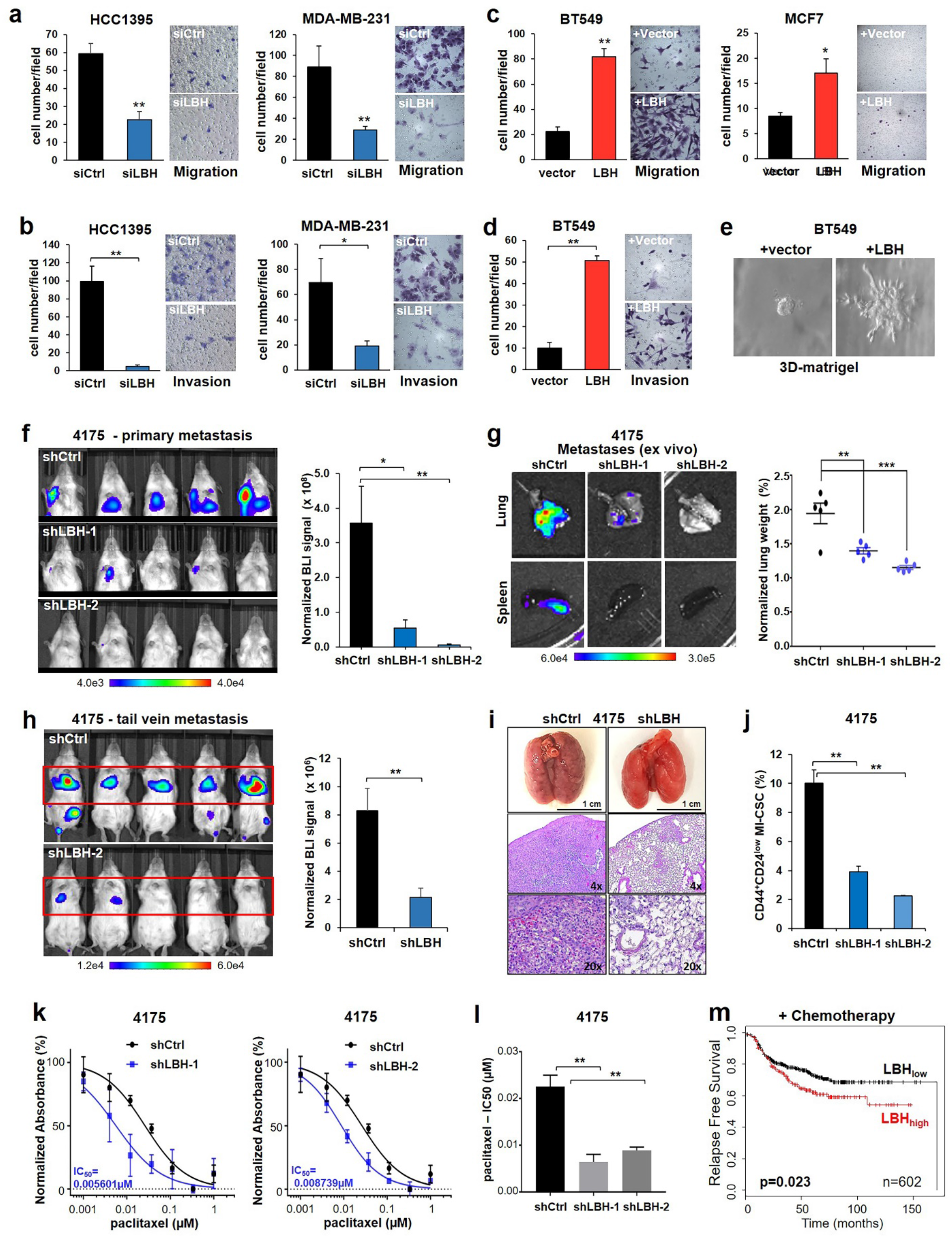
LBH promotes tumor cell motility, invasion, metastasis, and chemoresistance. (**a-d**) Boyden chamber assays, testing (**a,c**) cell migration, and (**b,d**) invasion in: (**a,b**) TNBC lines, HCC1395 and MDA-MB-231 transiently transfected with LBH siRNAs (siLBH) or scrambled siRNA (siCtrl); and (**c,d**) BT549 TNBC and MCF7 non-TNBC lines stably expressing LBH (+LBH) or vector control (+vector). Representative images of crystal violet-stained invasive cells (right panels). Two experiments with n = 3 replicates per group were performed. (**e**) Three-dimensional (3D) matrigel invasion assay (see Methods). Note, only in the presence of LBH (+LBH), BT549 form spheres invading extracellular matrix. (**f-g**) Orthotopic metastasis assay in 4175 TNBC cells stably expressing shLBH-1, shLBH-2, or non-target shCtrl. (**f**) Representative BLI images (left) and quantification of *in vivo* metastatic burden (right) 7 weeks after orthotopic transplantation of cells (10^5^). n = 6 mice per group. This experiment was performed twice. (**g**) *Ex vivo* BLI images of lung and spleen metastases (left) and quantification of lung metastatic burden (right) by lung weight measurements, normalized to the % of body weight. n=5 mice per group. (**h-i**) Tail vein metastasis assay using 10^5^ 4175 control (shCtrl) and LBH KD (shLBH-2) cells. (**h**) Representative BLI images (left) and quantification of lung metastatic burden (right) 5 weeks post engraftment. n=5 mice per group. (**i**) Representative images (top) and H&E-stained tissue sections (middle, bottom) of lungs at endpoint. Scale bar, 1 cm, or magnifications, as indicated. (**j**) FACS quantification of metastasis-initiating CD44^+^CD24^low^ CSC (MI-CSC). n = 4 per group. For FACS plots see Supplementary Fig. 2c. (**k,l**) 4175 cells were treated with DMSO or increasing concentrations of paclitaxel for 3 days (see Methods). (**k**) Standard dose response curves for 4175-shLBH1 and 4174-shLBH2 relative to 4174-shCtrl control (IC_50_ = 0.02609 μM), and (**l**) bar diagram comparing IC_50_ values between all three lines. n = 5 per group. Data represent means ± SEM. *P<0.05; **P<0.01; ***P<0.001. Two-tailed Student’s *t*-test (**a-d,h**) or One-way ANOVA (**f,g,j-l**) were used. (**m**) Kaplan-Meier analysis showing that top tier *LBH* expression (red) predicts reduced relapse-free patient survival after chemotherapy. Data were extracted from the Kaplan-Meier plotter dataset (http://kmplot.com/analysis). P=0.023 (log-rank test).

To examine the requirement for LBH in metastasis *in vivo*, MDA-MB-231-4175 cells expressing shCtrl or either of two LBH shRNAs (shLBH-1, shLBH-2) were injected orthotopically into NSG hosts and monitored for spontaneous metastasis formation. Loss of LBH drastically decreased the emergence of distant metastasis (p<0.01; n=6/group; Fig. 5f,g). While primary mammary tumors formed by 4175-shCtrl generated multi-organ metastases in lungs, spleen, and liver, 4175-shLBH cells formed either no metastasis or only lung micro-metastases, as determined by *ex vivo* bioluminescence imaging of organs (Fig. 5g; left). Lung weights of 4175-shLBH-1- and 4175-shLBH-2-injected mice were also significantly decreased compared to controls, reflecting their reduced lung metastatic burden (Fig. 5g; right). Moreover, when injected directly into the blood stream via tail vein, the shLBH lines also showed a markedly reduced ability to colonize the lung parenchyma (shown for shLBH-2; Fig. 5h,i). Notably, the number of metastasis-initiating CD44^+^CD24^low^ CSC (MI-CSC) was significantly reduced by >2.5 to 4-fold in LBH-depleted 4175 compared to 4175-shCtrl control cells (p<0.01; Fig. 5j), indicating that loss of metastatic potential upon LBH depletion resulted from reduced CSCs. Thus, LBH is required for both metastasis from primary tumors and for extravasation, and establishment of tumor metastases in this highly aggressive TNBC model.

Since breast cancer death results from recurrence of treatment resistant metastases, we also evaluated *LBH* and treatment resistance. Notably, LBH depletion in chemotherapy-resistant 4175 TNBC cells with two shRNAs markedly increased sensitivity of 4175 to paclitaxel (3-4-fold; *p*<0.01; Fig. 5k,l), a chemotherapy drug widely used for breast cancer. Conversely, ectopic expression of LBH in highly chemo-drug sensitive MCF7 cells resulted in paclitaxel resistance (data not shown). To evaluate the clinical significance of these findings, we performed Kaplan Meier analysis of published breast cancer gene expression data. Notably, in a cohort of 3,951 breast cancer patients, of which 602 were treated with neoadjuvant or adjuvant chemotherapy, high intra-tumoral *LBH* associated significantly with reduced relapse-free survival after chemotherapy (Fig. 5m). Thus, high *LBH* levels correlate with chemoresistance in patients, and LBH inhibition has the potential to restore chemo-drug response.

Collectively, these data uncover a novel role for LBH in malignant tumor progression, as a driver of CSC expansion, chemoresistance and aggressive metastatic TNBC recurrence.

### LBH activates mammary stem cell programs and represses E-cadherin adherence junction genes

To investigate the mechanisms underlying the CSC-promoting effects of LBH, we performed comparative global gene expression analysis by RNA-Seq in all four breast cancer cell models above. GSEA analysis of differentially expressed genes revealed a significant enrichment of stem cell-associated gene signatures (P<0.05; FDR q<0.25) by LBH (Fig. 6a,b; and data not shown). Notably, the MaSC gene signature from Pece et al. ^35^ was most strongly and significantly upregulated in LBH overexpressing lines (MCF7+LBH/NES=+1.9; q=0.011 and BT549+LBH/NES=+1.7; q=0.052), whereas it was downregulated in LBH knockdown lines (HCC1395-siLBH/NES=-3.17; q<0.001 and MDA-MB-231-siLBH/NES=-1.77; q=0.026) vs. respective controls (Fig. 6a,b). Thus, LBH activates stem cell gene expression programs.

**Figure 6:**
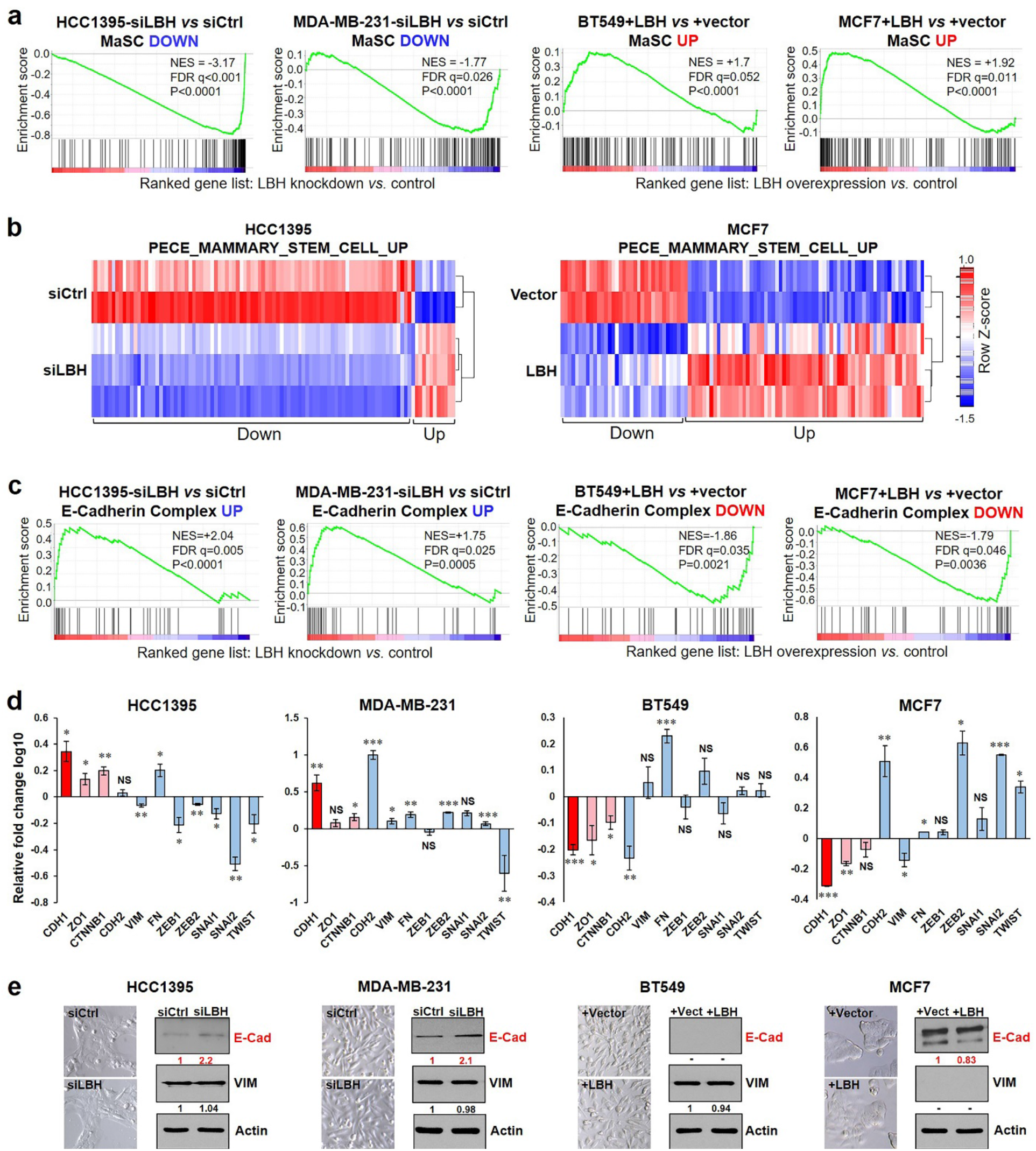
LBH activates mammary stem cell programs and represses E-cadherin adherence junction genes. (**a-c**) RNA-Seq analysis of HCC1395 and MDA-MB-231 TNBC cells three days post transient transfection with LBH-targeted siRNAs or scrambled siRNAs, and BT549 TNBC and MCF7 non-TNBC cells stably expressing LBH or a vector control (n=3/per group). (**a**) Gene set enrichment analysis (GSEA) shows the Pece_Mammary Stem cell (MaSC) signature ^35^ was most strongly enriched in a total of 8 common and significantly (P<0.05, FDR q<0.25) LBH-upregulated gene sets in the MSigDB C2:CPG data base. NES, Normalized Enrichment Score. FDR, False Discovery Rate. (**b**) Heatmaps for HCC1395 and MCF7 are shown. (**c**) Enrichment plot: the PID_E-Cadherin_Stabilization_Complex gene signature was most highly enriched among a total of 7 commonly and significantly (P<0.05, FDR q<0.25) LBH-downregulated genes in the MSigDB C2:CP data base. (**d**) qPCR quantification of mRNA expression levels of epithelial (red) and mesenchymal (blue) markers and transcription factors, normalized to *GAPDH*. Data represent means ± SEM (>n=3, Student *t*-test). NS, not significant. *P < 0.05; **P < 0.01; ***P <0.001. (**e**) Cell morphologies (left panels) and Western blot analyses of EMT marker protein expression (right panels). Fold changes in E-cadherin and Vimentin protein levels (normalized to ß-actin) were quantified by densitometry.

In contrast, gene profiles associated with E-cadherin complex stability and adherence junction (AJ) pathways were highly enriched (p<0.05; FDR<0.25) among top LBH-downregulated pathways (Fig. 6c; and data not shown), suggesting LBH represses epithelial cell differentiation. As downregulation of E-cadherin is also a hallmark of the Epithelial-Mesenchymal-Transition (EMT), a genetic program associated with metastasis and tumor cell dedifferentiation ^38,39^, we investigated EMT gene signatures in our RNA-Seq data sets, as well as EMT marker expression by qPCR and WB analysis. EMT gene signatures and expression of key EMT markers and EMT-TFs were not consistently changed, nor did we observe EMT-like changes in cell morphology, following LBH up- or downmodulation, except for E-cadherin (Fig. 6d,e; and data not shown). As observed by RNA-Seq, E-cadherin was significantly upregulated at both the RNA and protein levels by LBH loss in HCC1395 and MDA-MB-231 and downregulated by ectopic LBH in MCF7 and BT549 (Fig. 6d,e). From these studies, we conclude that LBH increases breast cancer cell motility, invasion, and metastasis by downregulating E-cadherin adherence junction genes.

### LBH is required for WNT-driven breast CSC expansion *in vivo*

Increasing evidence indicates that the oncogenic WNT/ß-catenin stem cell signaling pathway is hyperactivated in TNBC relative to other breast cancer types^3,4,16,40^, and required for breast CSC self-renewal/maintenance, metastasis, and chemoresistance ^40–43^. However, WNT downstream mechanisms in breast CSCs and TNBC carcinogenesis remain poorly understood. Our GSEA of LBH^high^ primary TNBC tumors above (Fig. 1d,e) and prior work ^16,17,20^ indicate a strong link between WNT and aberrant LBH expression in TNBC. We, therefore, asked if LBH, as a direct WNT/β-catenin target gene ^16^, plays a role in WNT-mediated breast CSC control.

To address this genetically, we tested the effects of *Lbh* knockout (KO) in a mouse model of WNT-driven TNBC, MMTV-Wnt1^Tg^ mice ^28,44^. These mice exhibit abnormal amplification of MaSCs in pre-neoplastic mammary glands ^45^ and enrichment of CSCs in tumors ^46^ due to the selfrenewal effects of WNT, which drive early tumor onset and malignant transformation in this *in vivo* model ^47^. In contrast, we previously showed that LBH KO in mice reduces normal MaSC frequency ^48^. MMTV-Wnt1^Tg^ mice were crossed with LBH KO mice (ROSA26[R26]-Cre;Lbh^-/-^) ^48^, and the SHIP-GFP line (Fig. 7a), which expresses GFP specifically in activated MaSCs ^49^, to allow *in vivo* stem cell analysis. Loss of LBH profoundly delayed tumor formation, with a median tumor-free survival of 50 weeks in MMTV-Wnt1+;SHIP+;Lbh^-/-^ KO compared to 10.5 weeks in MMTV-Wnt1^+^;SHIP+;Lbh^+/+^ wild type (WT) mice (p=0.0026; Fig. 7b). Notably, sphere formation of FACS-purified MaSCs (CD29^high^CD24^+^) from pre-neoplastic mammary glands (Fig. 7c), and the frequency of SHIP-GFP+ tumor stem cells in mammary tumors from LBH-deficient MMTV-Wnt1 mice were significantly and importantly reduced (Fig. 7d,e). Thus, LBH not only promotes CSCs in human breast cancer models but is also an essential effector of WNT-driven CSC expansion in mammary tumors *in vivo*.

**Figure 7:**
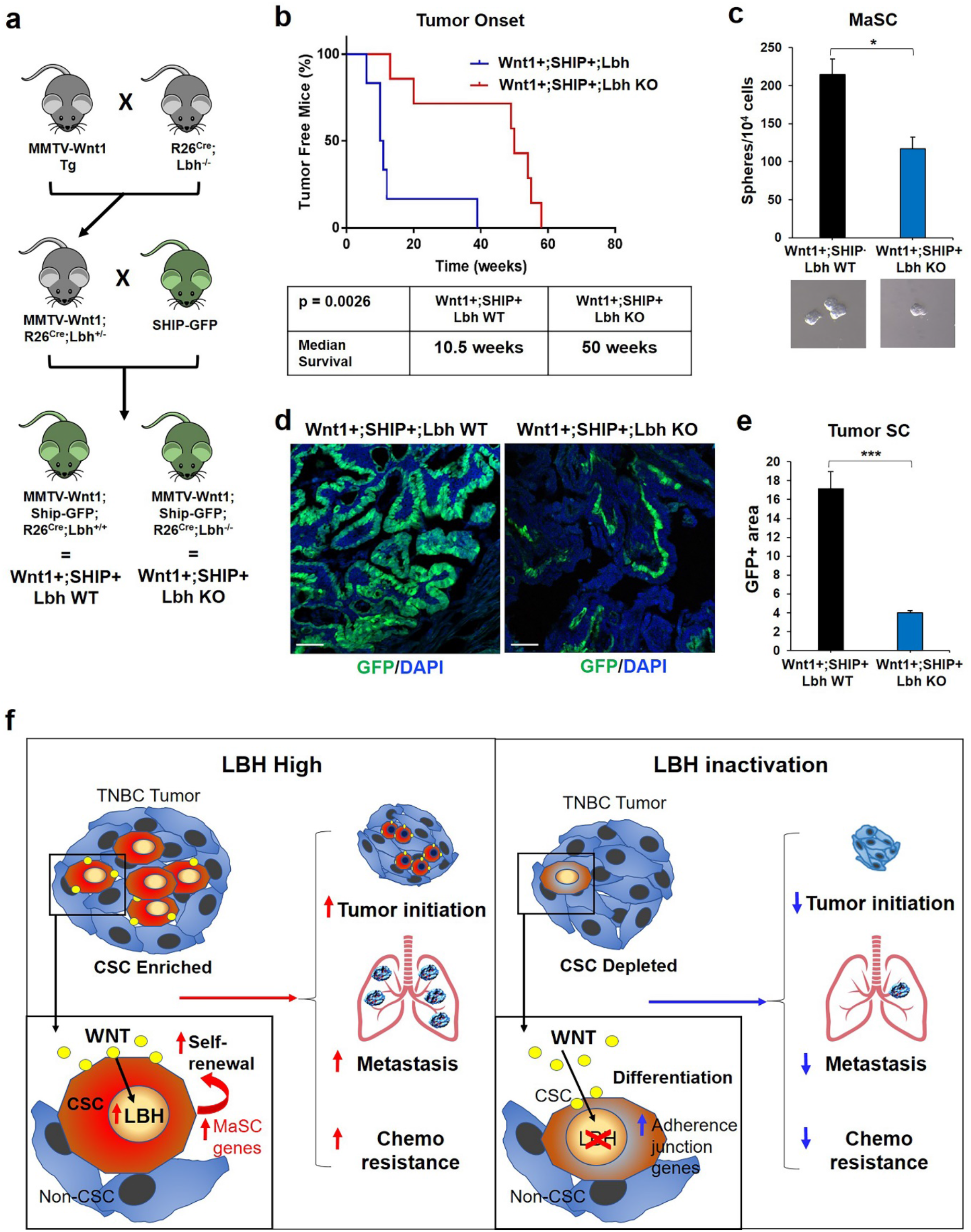
LBH is required for WNT-driven tumor stem cell expansion *in vivo*. (**a**) Experimental design: MMTV-Wnt1^Tg^ mice were crossed with ROSA26(R26)-Cre;Lbh-/- knockout (KO) mice and SHIP-GFP stem cell reporter mouse line. (**b**) Kaplan-Meier plot of tumor-free survival in MMTV-Wnt1;R26Cre;SHIP+;Lbh^+/+^ wild type (WT) (n=6) and MMTV-Wnt1; R26Cre;SHIP+;Lbh^-/-^ KO (n=7) female mice. Median survival data (Table, bottom). P=0.0026 (log rank test). (**c**) Sphere formation of FACS-purified CD29^high^CD24^+^ mammary stem cells (MaSC) from 12-week-old mammary glands (n=3 mice/group and triplicate samples). (**d**) Representative Immunofluorescence images and (**e**) quantification of SHIP-GFP+ tumor stem cells (SC) in tissue sections of mammary tumors from (b), using ImageJ software (>n=3 fields/section/group). Data represent means ± SEM. *P<0.05; ***P<0.001. Two-tailed Student’s *t*-test (**c,d**). (**f**) Schematic: Roles of WNT effector LBH in promoting breast CSC self-renewal, tumor initiation, metastasis and chemoresistance. LBH inhibition attenuates these carcinogenic processes by inducing CSC differentiation.

## DISCUSSION

Here we identified that LBH, a poorly characterized WNT target transcriptional regulator ^16^, is a critical breast CSC-intrinsic WNT effector that sustains high CSC abundance, metastasis, and chemoresistance in TNBC (Fig. 7f); and thus contributes to the aggressiveness and treatment resistance of these lethal cancers. Mechanistically, LBH activates mammary stem cell transcriptional programs, while it downregulates epithelial adherence junction genes. These data shed new light on the cell-intrinsic mechanisms governing TNBC aggressiveness, CSC transformation during tumor progression, as well as highlight LBH as a novel molecular target for CSC-specific cancer therapy.

While *LBH* gene expression has been shown previously to be elevated in TNBC-enriched molecular subtypes of breast cancer ^16,18^, we uniquely report the potential of LBH to stratify TNBC into a subset of stem-like cancers, independent of other molecular subclassifications. LBH was expressed in both basal-like and mesenchymal TNBC, and LBH^high^ TNBC tumors were enriched in stem cell pathways, most significantly WNT, consistent with *LBH* being a WNT/ß-catenin target gene ^16^. Importantly, we found LBH is specifically expressed in CD44^+^CD24^-/low^ breast CSCs, and its expression increases with the degree of cancer stemness. Thus, LBH may have unique utility as a biomarker to detect CSC-enriched breast cancers in the clinic.

Notably, silencing of LBH in multiple TNBC cell lines, and in primary patient-derived TNBC tumor cells reduced CD44^+^CD24^-/low^ CSC frequency and/or stem-like activity, while ectopic expression of LBH in BT549 and primary TNBC cells increased these CSC parameters. Importantly, depletion of LBH in highly malignant 4175 TNBC cells restored responsiveness to chemotherapy, and decreased TI-SC frequency, tumor growth, and metastasis *in vivo*, indicating LBH has potential as therapeutic target for anti-cancer stem cell therapy.

The CSC-promoting effect of LBH was not limited to TNBC; rather LBH may have a universal effect in promoting stemness during breast cancer progression. This notion is supported by our findings that ectopic LBH expression in the luminal ER+ MCF7 breast cancer model also increased the CD44^+^CD24^-/low^ CSC population, as well as self-renewal *in vitro* and tumor formation *in vivo*. Moreover, high intra-tumoral *LBH* expression correlated with increased metastasis, chemoresistance, and death in the entire patient population in different data sets.

We also investigated the potential causative role of LBH in WNT-mediated CSC expansion. Using a genetic mouse model for WNT-driven TNBC ^28^, MMTV-Wnt1^Tg^, and stem cell reporters, we show that LBH inactivation in MMTV-Wnt1^Tg^ mice significantly reduced MaSC activity at early stages and frequency of SHIP-GFP+ tumor stem cells at later stages, causing a profound delay in tumor onset. Thus, LBH not only promotes CSCs in human breast cancer, but is also an essential effector mediating WNT-induced CSC amplification *in vivo.*

During vertebrate embryonic development, *Lbh* is expressed in stem cell niches and organizer regions controlling tissue formation, i.e. pluripotent Xenopus blastomeres ^50^, chick dorsal mesoderm ^51^, and murine Apical Ectodermal Ridge (AER) ^13^, where we showed *Lbh* expression is controlled by WNT signaling ^16^. Moreover, developmental *Lbh* gain-of-function studies in mouse and chick have shown LBH promotes an undifferentiated, proliferating fetal progenitor state, while blocking cell differentiation ^14,52^. Yet, global *Lbh* KO in mice does not impair fetal development, vitality, or essential adult organ function ^48^. This indicates that LBH is not essential for stem cell function during embryogenesis or adult tissue homeostasis. However, LBH, which is expressed in mouse and human MaSCs after birth ^19,53^, is critically required for the rapid MaSC expansion that drives the extensive mammary tissue growth during puberty and pregnancy ^19^. Thus, LBH appears to be a unique vertebrate stem cell regulator needed only in situations requiring increased adult stem cell activity.

Increased stem cell function and tumor dedifferentiation are also hallmarks of cancer ^54^. It is therefore remarkable, that we found LBH is sufficient to confer stem-like traits on breast cancer cells, while it is critical for maintaining high CSC abundance in TNBC, and in WNT-driven mammary tumors *in vivo*. Despite the paramount importance of WNT as a CSC-promoting pathway ^55^, no WNT-targeted therapies are currently approved in the clinic, as inhibiting WNT signaling components often is cytotoxic, due to a requirement of WNT for normal tissue homeostasis, i.e. in the gut ^56^. Our work provides a novel rationale for targeting LBH to inhibit the CSC-promoting effects of WNT, with likely little side effects in patients. Future studies are needed to test the potential therapeutic benefit of LBH inhibition in diverse cancers driven by WNT.

*LBH* is also a direct TGFβ/SMAD3 target gene, however induction of *LBH* by TGFβ appears limited to rare claudin-low TNBC ^18^. Interestingly, in claudin-low TNBC lines, TGFβ promotes CSCs ^57^, and LBH knockdown studies in claudin-low MDA-MB-231 suggest that LBH is required for the CSC-promoting effect of TGFβ in this TNBC subtype ^18^. In contrast, in luminal MCF7 breast cancer cells, which we showed lack LBH, TGFβ acts as tumor suppressor and inhibits TI-SC activity ^57^. Thus, LBH may be required downstream of multiple CSC-inducing signaling pathways.

Our global transcriptomic analysis suggests that LBH promotes breast CSCs and metastasis by activating stem cell transcriptional programs, and by downregulating E-cadherin adherence junction genes ^58^, a key feature of epithelial differentiation. Unlike other factors with similar stemness- and metastasis-promoting properties, i.e. SNAIL, SLUG, TWIST, ZEB1/2 ^38,39^, however, LBH did not induce mesenchymal, fibroblastic traits, upregulate VIMENTIN, and EMT gene signatures were not consistently changed in the multiple LBH knockdown and overexpression breast cancer lines. Moreover, in contrast to a recent study in gastric cancer cell lines, suggesting LBH may promote tumor cell invasion by upregulating Integrin/Focal Adhesion Kinase (FAK) signaling ^21^, FAK signaling pathway genes were consistently downregulated by LBH in breast cancer lines (data not shown). A priority of future investigation will be to elucidate the precise mechanisms whereby LBH induces a metastatic CSC phenotype.

Collectively, our study calls attention to LBH as a novel breast CSC-intrinsic WNT effector that promotes TNBC aggressiveness and tumor progression by potently conferring stem-like/metastatic programs on breast cancer cells. The CSC/metastasis-promoting role of LBH, revealed herein, may have broad clinical significance, as LBH is also overexpressed in an increasing number of other aggressive cancer types ^21,22,59,60^. Thus, LBH warrant further investigation as a potential candidate for CSC-targeted therapy.

## MATERIALS AND METHODS

### Immunohistochemistry (IHC)

Breast cancer tissue microarrays (Biomax BR961, BRC962, BR1503e) and paraffin sections from 24 clinical breast cancer samples representing five different tumor subtype (luminal hormone receptor/HR+ - low grade; luminal HR^neg^ - high grade; TNBC-metaplastic; TNBC-medullary; and TNBC-atypical medullary) were either incubated overnight at 4°C with custom-made, affinity-purified anti-LBH antibody, as described ^19^, or stained with commercial anti-LBH antibody (Sigma) using an automated Leica IHC staining platform. LBH immunostaining was scored by two pathologists blinded to the identity of the specimens. Scores were given as the percentage of carcinoma cell nuclei staining positive, with an absolute intensity on a scale of 0-3 (0, none; +1, low; +2, moderate; +3, strong homogenous immunostaining). Tumors with LBH immunostaining scores of 0 and +1 in tumor cell nuclei were considered LBH^low^, whereas tumors with immune staining scores of +2 and +3 and >50% of carcinoma cell nuclei staining positive for LBH were considered LBH^high^.

### Bioinformatics Analysis of Public Data

Gene expression data for *LBH* in primary human breast cancers were obtained from the following published datasets: TCGA ^25^, METABRIC ^26^, Lehmann ^30^, NKI-295 ^61^, Hatzis ^62^ (GSE25066), and Creighton ^36^ (GSE7513). DNA copy number information was extracted from the normalized SNP6.0 data downloaded from the METABRIC data set ^63^. These cohorts were divided into TNBC and non-TNBC tumors according to their IHC-defined biomarker status. TCGA gene expression values are log2 transformed RSEM values and downloaded from UCSC Xena (https://xenabrowser.net). TCGA breast cancer subtypes were identified based on the gene expression using TNBCtype ^64^. LBH^high^ and LBH^low^ samples were defined relative to the median LBH expression level of TCGA samples of the selected TNBC subtypes (BL1 and BL2). Gene set enrichments in LBH^high^ vs. LBH^low^ TNBC tumors in the TCGA data set ^25^ was analyzed using Gene Set Enrichment Analysis (GSEA) ^65^. LBH^high^ and LBH^low^ patients were defined relative to the median LBH value of all patients in the NKI-295 data set. Kaplan-Meier analysis and log-rank test of survival difference between LBH^high^ and LBH^low^ patients were performed using the *survival* package from R (ver. 3.5.0). For meta-analysis of correlation with chemoresistance in breast cancer, we used the KM plotter http://kmplot.com/analysis). Patient samples were selected based on whether they received neoadjuvant or adjuvant chemotherapy and grouped for high or low mRNA expression of LBH. All percentiles between the lower and upper quartiles were computed, and the best performing threshold was used as cutoff.

### Cell lines

The breast cancer cell lines MCF7, T47D, ZR751, SKBR3, BT474, BT549, BT20, MDA-MB-361, MDA-MB-231, MDA-MB157, MDA-MB-436, MDA-MB-468, HCC1395, HCC1937, HCC1187, HCC1500, DU4474, and the normal-derived MCF10A cells were newly purchased from ATCC. The breast cancer cell lines, SUM52, SUM149, SUM 1315 were from Asterand. CAL51 was kindly provided by Dr. Chris Lord (ICR, London), and MDA-MB-231 derivative, 4175 ^31^, expressing luciferase by Dr. Joan Massague (Sloan Kettering Institute). Primary HMEC cells were from Lonza; and HMLE and HMLER cells from Dr. Priya Rai (University of Miami). All cell lines were maintained as recommended by the suppliers. Specifically, BT549, MDA-MB-231, 4175, MDA-MB-157 were grown in DMEM+10% FBS, Non-essential Amino Acids and Pen-Strep; HCC1395, HCC1187 in RPMI+10% FBS, Pen-Strep; and MCF7 in IMEM+10%FBS, Insulin and Pen-Strep in a 5% CO_2_ incubator at 37°C. Mycoplasma tests were routinely performed using MycoAlert Mycoplasma Detection Kit (Lonza). Stable LBH-overexpressing BT549 cell lines were generated by nucleofection of 2×10^6^ BT549 cells with 2 μg linearized pCDNA3 or pCDNA3+Lbh ^13^ and 1 μg pEGFP (Lonza) in Solution V (Lonza), using program A-023 on an Amaxa nucleofector. 48 hours post nucleofection, growth media containing 350 μg/ml G418 was applied to the cells to select for stably integrated transfectants. MCF7 cells were stably transfected with pCDNA3 or pCDNA3+Lbh ^13^ using Lipofectamine (Invitrogen), and after selection in 300 μg/ml G418 cells were tagged with luciferase by transduction with lentiviruses expressing pFU-Luc2-eGFP (Addgene).

### Small interfering RNA (siRNA) and short hairpin RNA (shRNA)-mediate gene knockdown

For RNAi studies, triplicate samples of cells were transiently transfected with 2 nM of synthetic siRNA specific for *LBH* or a scrambled control sequence (Dharmacon SmartPool) using Dharmafect #1 transfection reagent (Dharmacon) according to manufacturer’s protocol. Cells were incubated with siRNA containing media for 72 hours prior to splitting for other studies. For stable LBH knockdown, MDA-MB-231 (ATCC) and MDA-MB-231 variant, 4175 (J. Massague), were transduced with ready-made Mission shRNA lentiviral particles (Sigma) expressing four different *LBH*-specific shRNAs (TRCN0000107525-shLBH#1, TRCN0000107529-shLBH#2, TRCN0000107538-shLBH#3 and TRCN0000107533-shLBH#4) or a non-targeting control shRNA (Sigma, SHC002V) at MOI=5 and in the presence of 8 μg/ml polybrene, as in ^66^. Individual MDA-MB-231 and 4175 polyclonal cultures stably expressing LBH shRNA or control shRNA were obtained by selection in 2-5 μg/ml puromycin for > 10 days.

### Quantitative Real-time PCR (qPCR)

Total RNA was extracted from cells using Trizol reagent (Invitrogen), and reverse transcribed with M-MLV Reverse Transcriptase (Promega). qPCR was performed using SsoFast Evagreen PCR master Mix (BioRad) and a Bio-Rad CFX96 Thermal Cycler. mRNA expression was normalized to the expression of *GAPDH* using standard comparative Ct method. Sequences of qPCR primers upon request.

### Western Blot analysis

Cells were harvested in RIPA lysis buffer (Thermo Fisher Scientific, MA, USA) supplemented with protease and phosphatase inhibitor cocktails (Sigma-Aldrich, UK). Cell lysates were passed 5-8 times through a 26-gauge needle before centrifugation at high-speed. Cleared lysates were snap frozen until further use. 20-50 μg of total protein lysates were separated under reducing conditions (2.5% β-mercaptoethanol) by SDS-PAGE (SDS-Polyacrylamide Gel Electrophoresis) and transferred to nitrocellulose membrane (Amersham GE, UK) using Amersham ECL semi-dry blotter (GE). Membranes were incubated with primary antibodies to LBH (Sigma 1:1,000); OCT4 and NANOG (Novus 1:1,000); E-Cadherin (BD; 1:1,000); Vimentin (Sigma 1:1,000); or β-actin (Sigma, 1: 50,000) in TBST + 5% milk, followed by incubation with secondary HRP-coupled anti-rabbit or anti-mouse antibodies (Santa Cruz 1:10,000). Protein bands were detected using the West Femto Super Signal Kit (Thermo) on X-ray film, and quantified by densitometry and ImageJ analysis.

### Proliferation and Clonogenicity Assays

Cells were seeded in quadruplicates at 2 x 10^4^ cells per well of 24-well plates. Cell numbers were counted daily over 6 days. Data were normalized to values measured at day 0. For two-dimensional (2D) colony formation, cells were plated in triplicate at low density (100 - 500 cells per well) in 6-well plates and grow for 2 - 3 weeks with regular media changes. Cells were fixed in methanol and stained with 0.35% crystal violet solution. Colonies were counted manually or using Gelcount (Oxford Optronix). For threedimensional (3D) anchorage-independent soft agar growth assays, 0.6% Noble Agar (BD) was used as bottom layer in a 6-well plate and topped with a layer of 0.3% noble agar mixed with 5 x 10^3^ (MDA-MB-231, HCC1395, MCF7) or 2 x 10^4^ (BT549) cells in selective drug-containing growth medium. Triplicate samples of single cell suspensions were plated and grown for 21-28 days at 37°C in a 5% CO_2_ incubator, with addition of 100 μl of fresh growth media every 2 days. Colonies were stained with 0.05% crystal violet for > 2 hours, photographed, and colonies larger than 50 cells or 8 megapixels at 100% magnification were counted.

### Mammosphere Assays

Single cells were plated in triplicates on low-attachment 6-well plates (Corning) at a density of 2-5 x 10^3^ (human breast cancer lines) or 1×10^4^ (primary mouse mammary epithelial cells) cells per well in serum-free DMEM/F12 medium supplemented with 20 ng/ml EGF, 20 ng/ml bFGF, B27 supplement (Invitrogen) diluted 1:50, and 1 mg/ml penicillin/streptomycin, as described ^19^. Spheres were allowed to form for 5-14 days at 37°C in a 5% CO_2_ incubator and then quantified. For serial passaging, primary tumor spheres after 7 days in culture were collected in culture media by centrifugation at 450 g for 5 min, washed with PBS, trypsinized, and mechanically dissociated into a single cell suspension using a 21-gauge syringe. After washing in media containing 2% heat inactivated FBS (HI-FBS), cells were resuspended in PBS for counting and re-plating in secondary mammosphere cultures ^19^.

### Infection and sphere assays of primary breast cancer cells

Fresh tissues from a primary triple-negative breast cancer (TNBC1) resected from a patient after neoadjuvant chemotherapy was obtained from the Biospecimen Shared Resource, Sylvester Comprehensive Cancer Center, University of Miami. Tumor tissues were minced into small pieces and dissociated into single cells using human tumor tissue dissociation kit (Milteny #130-095-929) and Milteny gentle MACS tissue dissociator at 37°C for 1 hour according to the manufacturer’s protocol. Cells were treated with DNase I (StemCell Technologies #07900), washed with PBS, followed by RBC elimination using ACK Lysing solution (Thermo). Cells were washed again with PBS and passed through a 40-μm nylon strainer. Thereafter, cells were pelleted, resuspended in mammosphere medium and grown at low density (5,000 cells/ml) on ultralow-attachment plates (Corning) for 10-14 days to enrich for CSC. After four passages, TNBC1 spheres were dissociated into single cells by trypsinization for 5 min with 0.05% trypsin (Gibco, #25300-62). 1X trypsin inhibitor (1000X; 50mg/ml stock concentration; Sigma-Roche) was added to inactivate Trypsin, where after cells were washed two times with PBS and counted. 60,000 viable cells were resuspended in 5 ml mammosphere medium (see above) containing polybrene 1μg/ml (Sigma #107689). Cells were infected with 20 μl of concentrated lentiviruses expressing either LBH-specific Tet-shRNA vectors (KD1 and KD2) and non-target Tet-shRNA (Horizon Discovery), or pLVX TetOn-LBH expression vector (custom made) by spin transduction for 90 min at 1,000 g and 4°C, using a Thermo benchtop centrifuge equipped with a swing bucket rotor and 50ml tube adapters. After removing supernatants, cells were resuspended in fresh sphere medium and plated in triplicates -/+ 1 μg/ml Doxycycline (DOX) on 6-well ultralow attachment plates at 10,000 cells/well in 2 ml sphere medium. Sphere formation was assessed 10 days after DOX addition and growth at 37°C, 5% CO2 by counting and imaging. RNA was harvested from spheres to determine LBH knockdown and overexpression efficiencies by qPCR.

### Flow Cytometry, Cell Sorting

CD44-CD24 FACS analysis of human breast cancer cell lines was performed as in ^67^. Briefly, 1 x 10^6^ cells from subconfluent cultures in triplicates were resuspended in 100 μl ice cold PBS+2%FBS. Cells were immunostained with 20 μl each of anti-CD24-PE (BD Biosciences) and anti-CD44-APC (BD Biosciences) antibodies for 30 min on ice, washed with ice cold PBS+2% of heat-inactivated FBS (HI-FBS), and resuspended in 500 μl final volume of PBS+2% HI-FBS for analysis or sorting using LSR-II BD analyzer or FACS Arias, respectively. Data were analyzed using FlowJo software.

Primary mammary epithelial cells (MECs) were isolated from 12-week-old female mice, as previously described ^19^. Cells were blocked for 10 min in ice-cold PBS+2% heat-inactivated FBS (HI-FBS) containing anti-CD16/CD32 (BD Biosciences) and rat-γ-globulin (Jackson ImmunoResearch) antibodies. Cells were then immunostained for 30 min with APC-conjugated CD45, CD31 and TER119 antibodies (BD Biosciences) specific to Lineage (Lin) markers in combination with anti-CD24-PE and anti-CD29-FITC antibodies (BD Biosciences). Labeled cells were washed with ice-cold PBS+2% HI-FBS, incubated for 30 min with Streptavidin-APC (Invitrogen) and violet dead cell marker (Invitrogen) to exclude Lin+ and dead cells, filtered through a 40 μm filter (BD Falcon), and sorted using a FACS Aria-II (BD Biosciences). Sorted mammary stem cell (MaSC)-enriched CD29^high^CD24^+^ cells were then plated as single cell suspension in triplicates in low attachment 6-well plates at 10^4^ cells per well for mammosphere assays (see above).

### Cell migration/invasion assays

Boyden chamber Transwell migration and invasion assays were performed as in ^66^. For 3D Matrigel colony invasion assays, single cell suspensions of 2.5 x 10^3^ BT549 cells in 100 μl of complete growth media mixed with ice cold Matrigel (BD) (1:1) were plated in triplicate on 96-well plates. Plates were incubated at 5% CO_2_, 37°C for 30 min to allow the Matrigel to solidify, where after 100 μl of complete media + 200 μg/ml G418 was added to each well. The culture media was changed every 2 days. Ten to 14 days after plating, pictures were taken under bright field at 20X magnification, using Leica DMIL inverted microscope.

### Chemotherapy drug studies

MDA-MB-231-4175 (2.5 x 10^3^ cells per well) were seeded on 96-well plates in triplicates in complete growth media. After 24 hours, media was replaced with media containing paclitaxel (Sigma) at different concentrations (1 μM; 0.33 μM; 0.11 μM; 0.037 μM; 0.01 μM; 0.004 μM and, 0.001 μM). Cells were grown for an additional 72 hours, after which cell viability was quantified using CellTiter 96 AQ_ueous_ One Cell Proliferation Assay (Promega). IC_50_ concentrations were calculated by standard dose response curve method using Graphpad Prism software.

### Mouse Studies

For *in vivo* tumor xenograft studies, 5-week-old NOD-SCID IL2Rgamma^null^ (NSG) female mice [Stock No. 005557] were purchased from the Jackson Laboratories (Bar Harbor, ME). MDA-MB-231-4175 cells (10^5^), suspended in 100 μl PBS containing 50% Matrigel (BD), were injected unilaterally, or contralaterally at limiting dilutions (10^4^/10^3^/10^2^), into the inguinal mammary fat pads of 5-7-week-old female NSG mice, n=6-12 mice/group. For MCF7 tumor studies, female NSG mice were oophorectomized at 4 weeks-of-age and at 6 weeks-of-age received a 0.25 mg/90-day-release estrogen pellet, followed by unilateral, orthotopic injection of 2×10^6^ MCF7-Luc+LBH cells or control MCF7-Luc+vector cells per mouse (n=6-9 mice/group). For limiting dilution experiments (5×10^5^/5×10^4^/5×10^3^) cells were injected contralaterally into the 4^th^ mammary gland fat pad of NSG mice transplanted with a 0.18mg/90-day-release estrogen pellet. Primary tumor growth was quantified weekly by caliper measurement and Bioluminescence (BLI) Imaging using IVIS Spectrum *In Vivo* Imaging System (Perkin Elmer).

For orthotopic metastasis assays, primary tumors formed from orthotopic injection of 10^5^ 4175 cells (see above) were resected when they reached a size of 1 cm^3^. Mice were monitored weekly by BLI imaging to detect metastasis formation. For tail vein metastasis assays, 10^5^ 4175 tumor cells suspended in 100 μl PBS were injected intravenously (n=8 mice/group). At protocol-defined endpoints, primary tumors, lungs and other organs (spleen, liver) were dissected, weighed, and subjected to *ex vivo* BLI analysis for the detection of metastases and histopathological analysis.

For *in vivo* stem cell analysis in genetic mouse models, MMTV-Wnt1 transgenic (Tg) mice [B6SJL-Tg(Wnt1)1Hev/J; Stock No. 002870; Jackson Laboratories] were interbred with ROSA26(R26)Cre;Lbh^ΔE2/ΔE2^ knockout (KO) mice ^48^ and SHIP-GFP transgenic mice [B6.Cg-Tg(Inpp5d-EGFP)DLrr/CprJ; Stock No. 024808; Jackson Laboratories] ^49^. All experiments and procedures involving mice were approved by the Institutional Animal Care and Use Committee (IACUC) of the University of Miami in accordance with the National Institutes of Health Guide for the Care and Use of Laboratory Animals.

### In situ GFP analysis

Mammary tumor tissues from MMTV-Wnt1Tg;R26-Cre;SHIP-GFP+;LBH^+/+^ wild type (WT) and MMTV-Wnt1Tg;R26-Cre;SHIP-GFP+;LBH^-/-^ KO female mice were fixed in 2% paraformaldehyde for 3 hours, immersed in 30% sucrose overnight at 4°C, embedded in O.C.T, snap frozen on dry ice, and sectioned (10 μm). Tissue cryo-sections were mounted in Slowfade Gold Mounting Media with DAPI counterstain (Thermo). Endogenous GFP expression was evaluated by Confocal Microscopy on a Leica SP5 Inverted Confocal Microscope and quantified using ImageJ software.

### RNA-Seq Gene Expression Analysis

RNA from triplicate samples was extracted using Trizol (Invitrogen) and quantified by Agilent 2100 Bioanalyzer. RNA-Seq, including mRNA enrichment, library preparation, sequencing and data analysis, was performed by Novagen Inc. (California, USA). mRNA was enriched from 1 ug of total RNA starting material using Illumina TruSeq RNA Sample Prep Kit. RNA libraries were then prepared using NEBNext Ultra RNA-Seq library prep kit (New England Biolabs) followed by high-throughput sequencing on Illumina HiSeq 200. Between 38 and 60 million reads were obtained from each sample. Illumina Casava1.7 software was used for base calling. Raw sequence paired-ended data in FASTQ format were assessed for quality with FastQC (v11.5). Trimmomatic (ver.0.32) was used to remove adapters, Illumina-platform specific sequences, and low-quality reads. Reads were aligned to reference genome Homo sapiens GRCh37/hg19 (NCBI/UCSC/Ensembl) using STAR (v2.5). HTSeq v0.6.1 was used to count the read numbers mapped of each gene. Differential expression analysis for comparisons of two conditions (-/+LBH) per cell line (three biological replicates per group) with respect to their controls was performed using the DESeq2 R package (2_1.6.3). The resulting P-values were adjusted using the Benjamini and Hochberg’s approach for controlling the False Discovery Rate (FDR). Genes with FDR < 0.05 were assigned as differentially expressed. GSEA analysis was conducted using the Wald statistic output from DESeq2 results and permutation by gene set for MSigDB v7.1 (https://www.gsea-msigdb.org/gsea/msigdb/index.jsp) gene sets in the C2:CP, C2:CPG, C6, and Hallmark data sets ^65^. Only enriched gene sets commonly up-or down-regulated by LBH in all four cell lines (HCC1395, MDA-MB-231, BT549, MCF7) with p-values <0.05 and FDR q<0.25 were considered. The RNA-Seq data was deposited in the GEO database with accession number GSE151206.

### Statistical Analyses

Statistical analyses were performed using Microsoft Excel, GraphPad Prism Software, or statistical software package R (version 3.3.1). Each experiment was repeated three times or more and data are presented as mean ± standard deviation (SD) or standard error of the mean (SEM) unless otherwise noted. Data were analyzed by unpaired two-tailed Student’s *t-*test, or analysis of variance (ANOVA) for more than two group comparison. Chi-square test was used to compare categorical clinical variables with LBH protein expression status (low vs. high) shown in Table 1. One sample proportion test was used to examine whether proportion of overall high LBH protein expression among IDC is 50%. Gene expression and copy number differences in clinical samples were evaluated by two-sided Mann-Whitney U-test. The log-rank test was used for Kaplan-Meier survival analysis. Results with *p*<0.05 were considered statistically significant.

## Author Contributions

K. J.B. conceived the study. K.G. and K.J.B. designed the experiments. K.G., K.A-B., M.E.R., L. E.L, B.W., and K.J.B. performed the experiments. S.H. and R.Q. performed the MCF7 *in vivo* Xenograft experiments. D.J.A. helped with FACS analysis. K.G., K.A.-B., S.H., M.E.R. and L.E.L. analyzed the data. M.N. provided FFPE slides. M.N. and M.K. reviewed and scored the clincial IHC slides. D.K., Y.B., Z.G., S.X.C., and A.H.S. performed the statistical/bioinformatics data analysis. S.B.K. provided TNBC surgical specimen. C.J. and J.M.S. provided scientific input. J.M.S. edited the paper. K.G. and K.J.B. prepared the figures. K.J.B., K.G. and M.E.R wrote the paper.

## Acknowledgements

We thank Drs. Joan Massague for providing 4175 cells; Drs. Merce Jorda and Jennifer Chapman for paraffin sections and staining; Drs. Youping Sun and Pingping Chen for technical assistance; and Drs. Priya Rai and David Robbins for comments on the manuscript. We thank the Biological Biospecimen Shared Resources of the Sylvester Comprehensive Cancer Center at the University of Miami Miller Medical School for providing TNBC specimen. We also thank the Analytic Imaging, Cancer Modeling and Flow Cytometry Shared Resources for high resolution imaging, BLI analysis, or FACS analysis and cell sorting respectively. This work was supported by NIH/NIGMS Grant R01GM113256 (K.J.B), the Department of Defense (DoD)/Breast Cancer Research Program (BCRP) Breakthrough Award W81XWH-19-1-0255 (K.J.B); DoD/BCRP Pre-doctoral Fellowship W81XWH081053 (M.E.R.); and funding support from the Sylvester Comprehensive Cancer Center (K.J.B).

