## Supplemental data for "LBH is a cancer stem cell- and metastasis-promoting oncogene essential for WNT stem cell function in breast cancer"

### SUPPLEMENTARY INFORMATION

### SUPPLEMENTARY FIGURES

#### Supplementary Figure 1

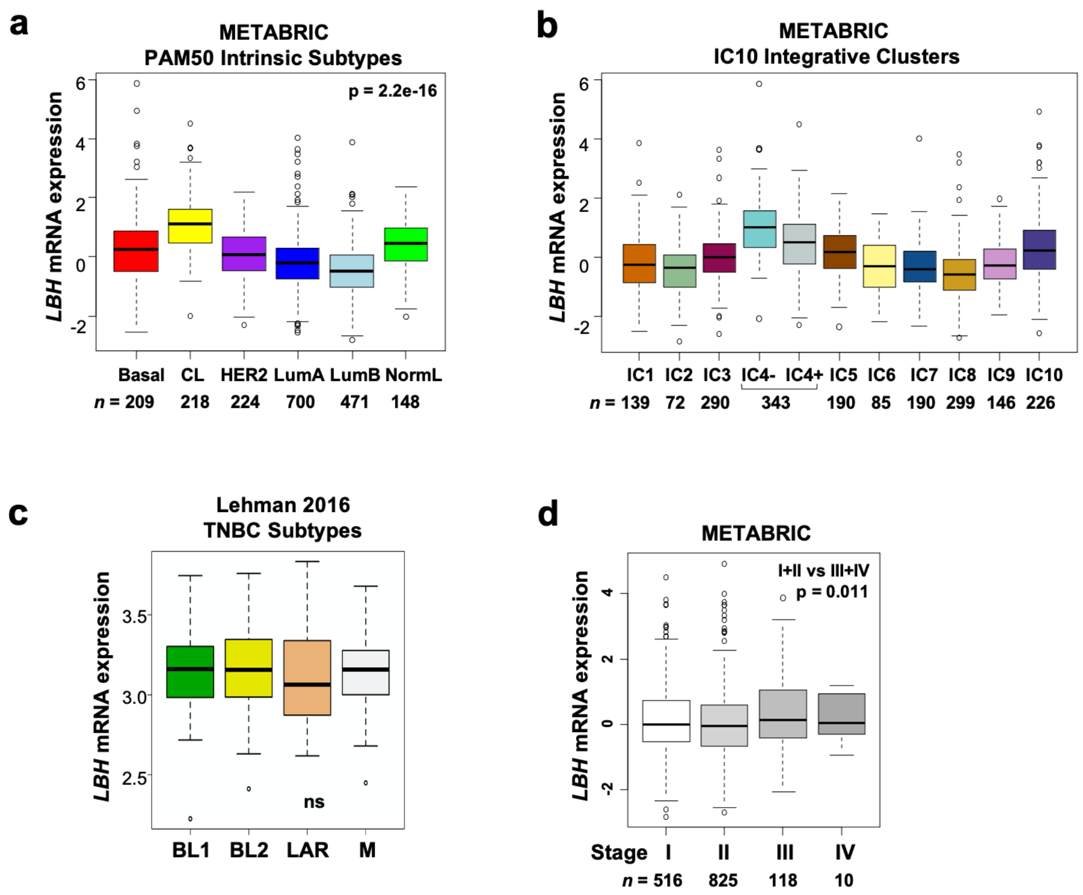


**Supplementary Fig. 1: LBH expression in primary human breast cancers grouped into molecular and prognostic subtypes (related to Main Fig. 1).** Gene profiling expression data for *LBH* classified by: (**a**) PAM50 intrinsic molecular subtypes [70]; and (**b**) IC10 Integrative Subtypes in 1,980 tumors from the METABRIC dataset [39]. All differences are significant p<0.05. (**c**) *LBH* expression in molecular subgroups of TNBC from the updated Lehmann data set [43], with highest *LBH* levels in basal-like 1 and 2 (BL1, BL2), mesenchymal (M) TNBC subtypes, and lowest levels in the luminal androgen receptor positive (LAR) TNBC subgroup. P-values by Mann-Whitney U test as indicated. ns, Not significant. (**d**) Significant correlation of *LBH* expression with advanced disease stage (III and IV; p=0.011). Number (n) of patients per group as indicated.

**Supplementary Figure 2**

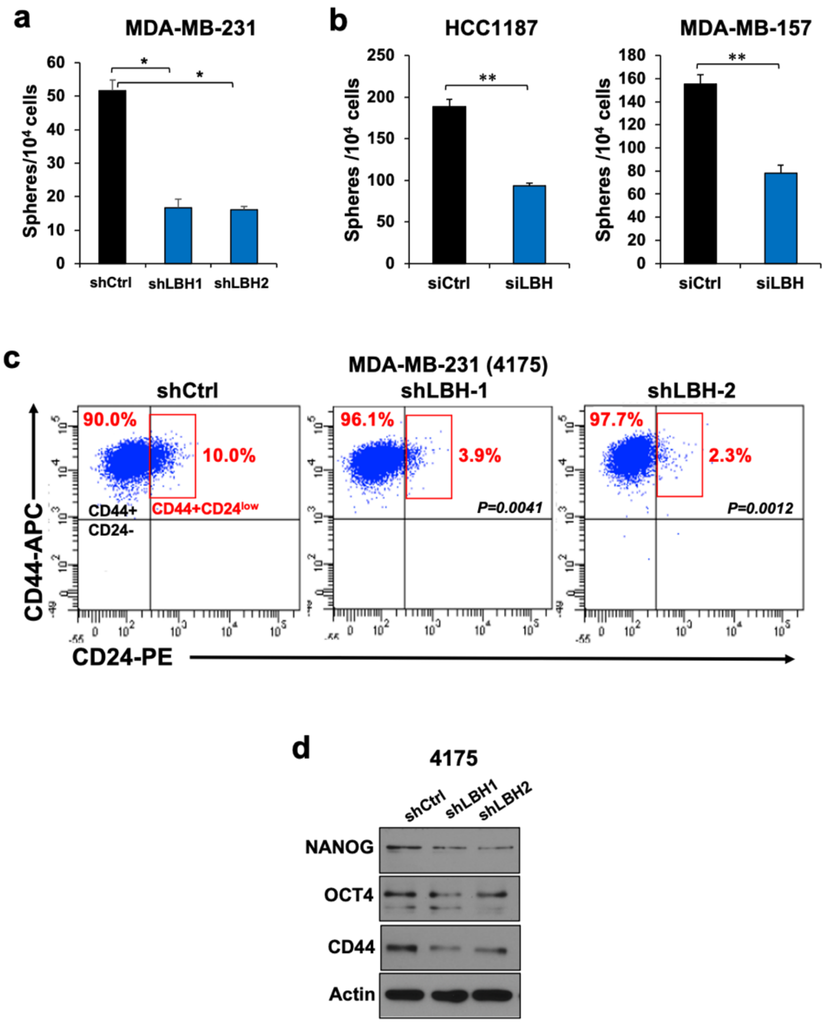


**Supplementary Fig. 2: LBH promotes breast cancer stemness (related to Main Fig. 4a-g).** (**a,b**) Tumor sphere formation assays in TNBC lines of different histopathological subtypes: (**a**) mesenchymal MDA-MB-231 stably transduced with shCtrl and shLBH-1, shLBH-2; and (**b**) basal A subtype HCC1187 and aggressive metaplastic MDA-MB-157 TNBC cells transiently transfected with non-targeted siRNA control (siCtrl) or a pool of 4 independent LBH-targeted siRNAs (siLBH). All data represent mean ± SEM (n=3, Student *t*-test; $P < 0.05$). (**c**) CD44-CD24 FACS profiles of the MDA-MB-231 sister line, 4175, stably transduced with shCtrl and two independent LBH-targeted shRNAs (shLBH-1, shLBH-2). Percentages of different CSC-enriched populations ($CD44^{+}CD24^{-}$ and $CD44^{+}CD24^{low}$) as indicated. Quantification of metastasis-initiating $CD44^{+}CD24^{low}$ CSCs (red box) is shown in main Figure 5j. (**d**) Western blot analysis of stem cell markers, NANOG, OCT4, CD44 and loading control, β-actin, in the same cells as in c.

**Supplementary Figure 3**

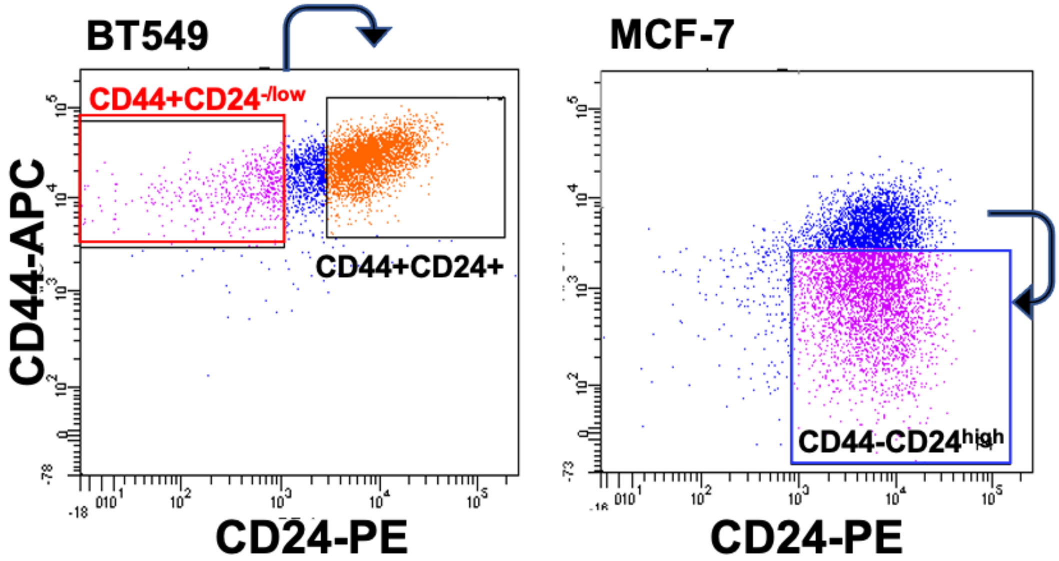


**Supplementary Fig. 3 (related to Main Fig. 5j):** CD44-CD24 FACS profiles of BT549 and MCF7 cells showing the distribution of breast CSC ($CD44^{+}CD24^{-/low}$; red rectangle) and non-CSC ($CD44^{-}CD24^{high}$ – blue rectangles) populations that were sorted for the qPCR gene expression analysis show in the main Figure 3b. Blue arrows indicate increasing degree of tumor cell differentiation.

**Supplementary Figure 4**

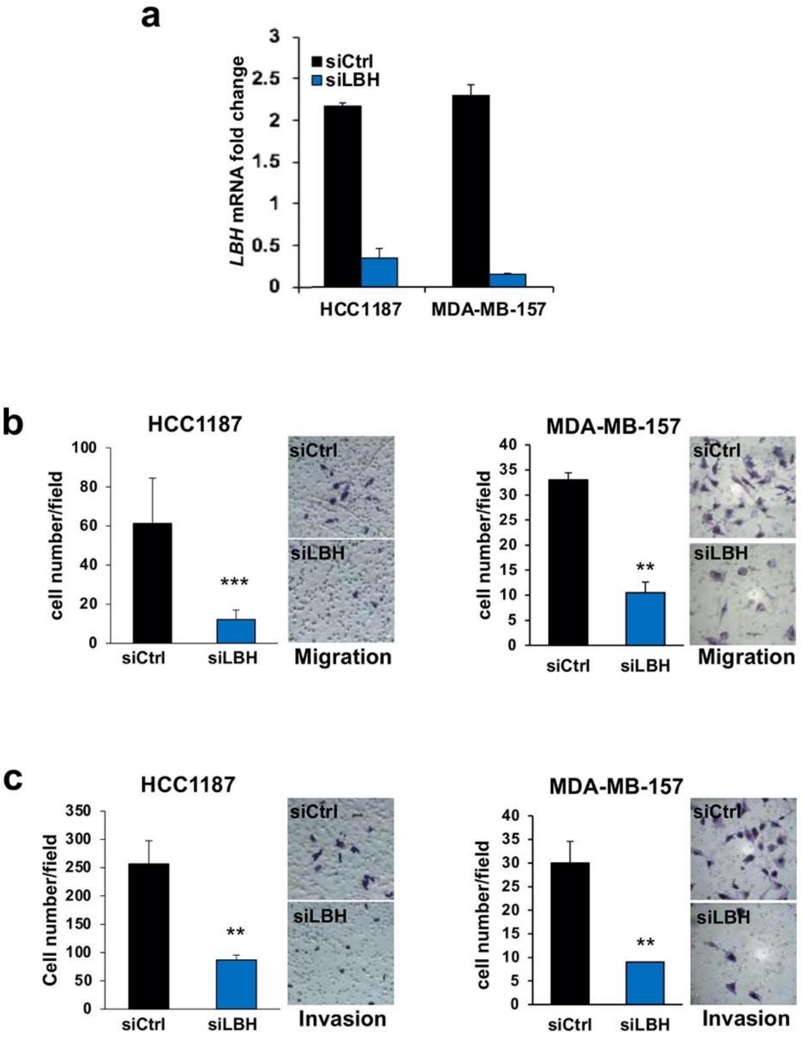


**Supplementary Figure 4: LBH induces TNBC cell motility and invasion (related to Main Figure 5a,c)**. HCC1187 (basal A) and metaplastic MDA-MB-157 TNBC cells were transiently transfected with non-target siRNA control (siCtrl) and LBH-targeted siRNAs (siLBH). (**a**) qPCR analysis of *LBH* normalized to *GAPDH* 3 days after siRNA transfection shows efficient LBH knockdown in siLBH transfected cells. Data represent means ± SEM (n=3, Student *t*-test; $P < 0.05$). (**b**) *In vitro* Transwell migration, and (**c**) Boyden Chamber Matrigel Invasion assays 4 days after transient transfection of HCC1187 and MDA-MB-157 TNBC cells with LBH-specific siRNAs (siLBH) or scrambled siRNA (siCtrl). Data represent means ± S.D. (n>3; Two-tailed unpaired *t*-test). *P*-values: $^*P < 0.05$; $^{**}P < 0.01$. Representative images of crystal violet-stained migratory tumor cells are shown on the right.

### SUPPLEMENTARY TABLES

**Supplemental Table 1: Cancer stem cell populations (%) in various breast cancer cell lines**

| Cell line | Tumor type | Tissue Source | Cell type classification | LBH expression | $CD44^{+}CD24^{-/low}$ |
|---|---|---|---|---|---|
| **HCC-1395** | DC | Primary | Basal/TN | +++ | 86.3 ± 4.42 |
| **MDA-MB-231** | AC | Pleural Effusion | Mesenchymal stem-like/TN | +++ | 96.8 ± 0.24 |
| **BT549** | IDAC | Primary | Mesenchymal/ TN | (+) | 11.8 ± 1.31 |
| **MCF7** | IDAC | Pleural Effusion | Luminal/ER+ | - | 0.47 ± 0.31 |

*TN* triple-negative (ER-/PR-/HER2-); *AC* adenocarcinoma, *DC* ductal carcinoma, *IDAC* infiltrating ductal adenocarcinoma
